# A screen for modulation of nucleocapsid protein condensation identifies small molecules with anti-coronavirus activity

**DOI:** 10.1101/2022.12.05.519191

**Authors:** Rui Tong Quek, Kierra S. Hardy, Stephen G. Walker, Dan T. Nguyen, Taciani de Almeida Magalhães, Adrian Salic, Sujatha M. Gopalakrishnan, Pamela A. Silver, Timothy J. Mitchison

## Abstract

Biomolecular condensates formed by liquid-liquid phase separation have been implicated in multiple diseases. Modulation of condensate dynamics by small molecules has therapeutic potential, but so far, few condensate modulators have been disclosed. The SARS-CoV-2 nucleocapsid (N) protein forms phase separated condensates that are hypothesized to play critical roles in viral replication, transcription and packaging, suggesting that N condensation modulators might have anti-coronavirus activity across multiple strains and species. Here, we show that N proteins from all seven human coronaviruses (HCoVs) vary in their tendency to undergo phase separation when expressed in human lung epithelial cells. We developed a cell-based high-content screening platform and identified small molecules that both promote and inhibit condensation of SARS-CoV-2 N. Interestingly, these host-targeted small molecules exhibited condensate-modulatory effects across all HCoV Ns. Some have also been reported to exhibit antiviral activity against SARS-CoV-2, HCoV-OC43 and HCoV-229E viral infections in cell culture. Our work reveals that the assembly dynamics of N condensates can be regulated by small molecules with therapeutic potential. Our approach allows for screening based on viral genome sequences alone and might enable rapid paths to drug discovery with value for confronting future pandemics.

## Introduction

Biomolecular condensates are membraneless organelles formed by liquid-liquid phase separation of specific RNAs and/or proteins, resulting in their local concentration in a liquid-like compartment distinct in constituents from the surrounding cytoplasm or nucleoplasm (1–3). Such biomolecular condensates have been implicated in the formation of signaling complexes, processing bodies, stress granules and germline bodies (1), where they facilitate the segregation and concentration of factors involved in various cellular processes. The material properties of condensates are tailored to their functions; dynamic condensates with mobile constituents enhance biochemical reactions that involve molecular turnover, whereas more glass-like or solid condensates promote stiffness for structural support (4).

Phase separation and the formation of liquid condensates known as viroplasms or inclusion bodies have been observed in large groups of viruses such as the *Mononegavirales* order of non-segmented negative-strand RNA viruses (5–16) and the *Reoviridae* family of double-stranded RNA viruses (17–19). The formation of viroplasms is induced by viral proteins and RNAs expressed during infection and they serve as organizational hubs for concentration of viral or host factors involved in viral entry, replication, virion assembly and/or packaging (20). The SARS-CoV-2 nucleocapsid (N) protein drives virion packaging through RNA-binding and enhances viral transcription and replication at replication and transcription complexes (RTCs) (21,22). Recent observations that the SARS-CoV-2 N protein forms liquid condensates (21,23–29) has raised the possibility that these N condensates may also behave as dynamic viroplasms. However, whether condensate assembly is a conserved property of HCoV Ns has not been examined.

Macromolecular phase separation is often driven by unstructured regions of proteins, which makes condensates unconventional for targeting by small molecule drugs. Nevertheless, recent studies successfully identified small molecules that modulate phase transitions of proteins involved in ALS (30,31) and respiratory syncytial virus infection (32). These studies, as well the recent founding of condensate-focused biotechnology companies, has led to an explosion of interest in targeting condensates for drug discovery, but to date, few active molecules have been disclosed (33–35). In principle, small molecules could achieve therapeutic activity by inhibiting the assembly of cytotoxic condensates (30,31), or by promoting condensation, leading to hardening and cessation of essential dynamics (32). Given the ongoing need for antivirals to confront the COVID-19 pandemic, and the likelihood that similar pandemics will emerge in the future, we focused on identifying small molecules that perturb SARS-CoV-2 N condensation, with the hope that some might exhibit broad-spectrum anti-viral activity. We developed a cell-based high-content screening platform to identify small molecules that either promote or inhibit N condensation and identified small molecules with condensate-modulating activity. Our results suggest it may be possible to discover drug-like small molecules that promote and inhibit condensation of many proteins and RNAs, which will open new paths to drug discovery.

## Results

### HCoV N condensates are polyIC-inducible and exhibit varied material properties

To investigate the condensation behavior of HCoV N proteins, we stably expressed each of the seven HCoV N proteins fused to a C-terminal EGFP in A549 cells (human lung cancer derived) (Figure 1A). Western blots confirmed that the expression levels of each of the seven N proteins were similar among the stably expressing A549 cells (Supplementary Figure 1A). Under control conditions, SARS-CoV, SARS-CoV-2, HCoV-OC43 and MERS-CoV N showed diffuse cytoplasmic localization, while HCoV-229E, HCoV-NL63 and HCoV-HKU1 N formed spherical condensates (henceforth referred to as ‘constitutive’ condensates) of varying numbers (Figures 1A-B). Thus, the tendency to phase separate and condense varies between N species under these conditions.

**FIGURE 1.**
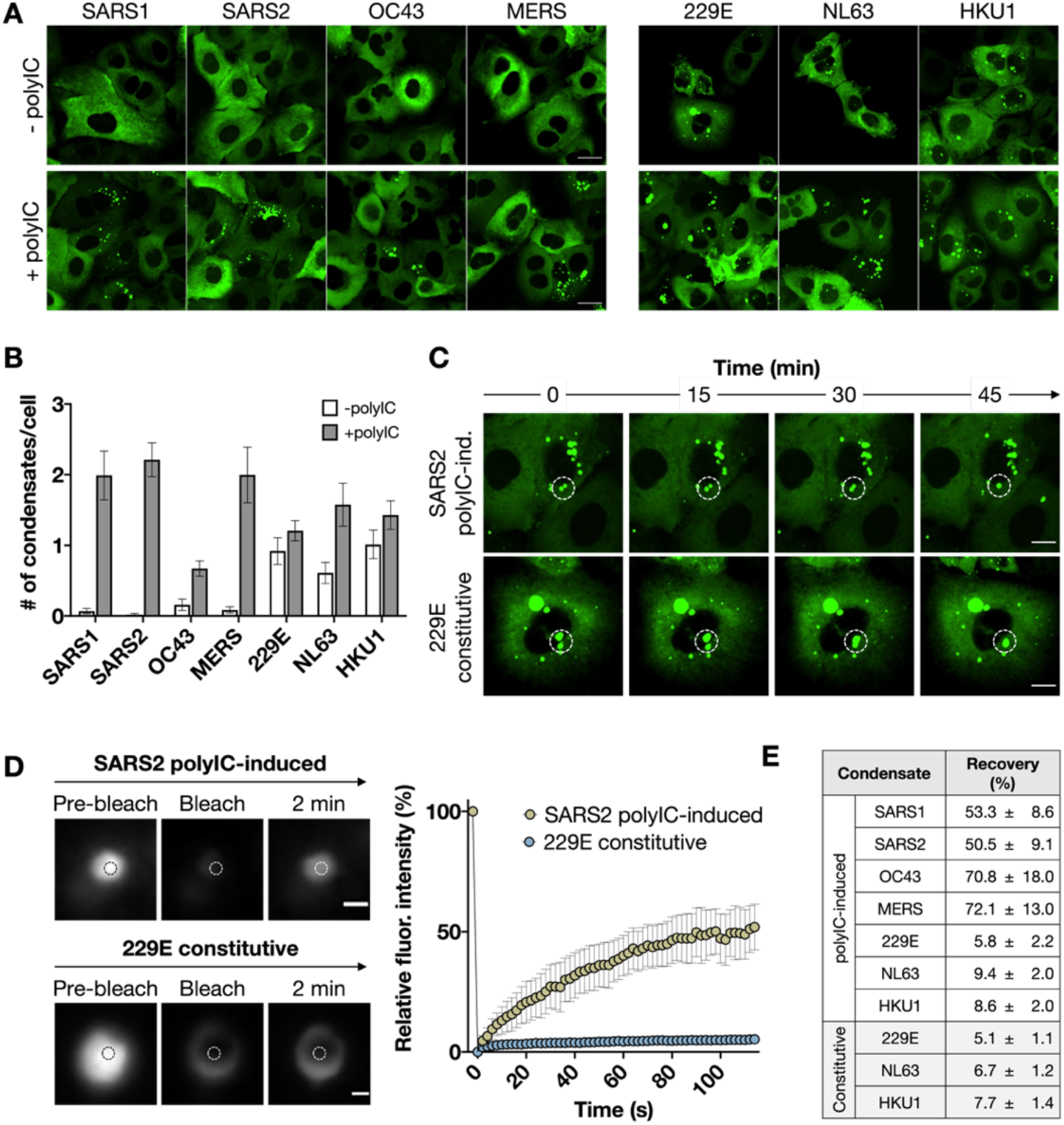
HCoV N proteins phase separate in a polyIC-inducible manner. (A) Fluorescence images of A549 cells stably expressing the N protein from seven HCoVs. For all seven cell lines, N condensates can be induced with 1μg/mL transfected polyIC for 7h. Images taken at 40x air magnification, confocal. Scale bar, 10 μm. SARS1, SARS-CoV; SARS2, SARS-CoV-2; OC43, HCoV-OC43; 229E, HCoV-229E; NL63, HCoV-NL63; HKU1, HCoV-HKU1; MERS, MERS-CoV. (B) Quantification of number of N condensates per cell for each of the seven cell lines. Data indicates mean ± standard deviation of duplicate experiments. (C) Time-lapse imaging showing polyIC-induced SARS-CoV-2 and constitutive HCoV-229E N condensate fusion events over time. Dotted circles highlight a representative fusion event. Images taken at 40x air magnification, confocal. Scale bar, 10 μm. (D) FRAP analyses of SARS-CoV-2 polyIC-induced and HCoV-229E constitutive N condensates. Images are single N condensates representative of each condition. Mean fluorescence intensity plot illustrates FRAP results for n = 7 condensates per condition. Images taken at 60x oil magnification, widefield. Scale bar, 1 μm. (E) Final fluorescence recovery percentage 2 min post-bleach. Data indicates mean ± standard deviation of all seven condensates.

Upon transfection of low molecular weight polyinosinic-polycytidylic acid (polyIC), a synthetic analog of dsRNA that mimics viral genome replication intermediates and triggers innate immune pathways, N condensates formed across all seven HCoV N cell lines (Figures 1A-B). For the species that exhibited constitutive condensation, the number of condensates increased upon polyIC transfection (Figures 1A-B). Addition of polyIC to an A549 cell line stably expressing EGFP alone did not result in condensate formation, confirming the essentiality of N for polyIC-induced condensate formation (Supplementary Figure 1B). Across all seven Ns, both constitutive and polyIC-induced condensates exhibit flow and fusion/coalescence over time (Figure 1C; Supplementary Figure 1C), which are behaviors consistent with liquid-liquid phase separation. We also probed the dynamics of the various N condensates by monitoring fluorescence recovery after photobleaching (FRAP). PolyIC-induced SARS-CoV/SARS-CoV-2/HCoV-OC43/MERS-CoV N condensates were much more dynamic than constitutive HCoV-229E/HCoV-NL63/HCoV-HKU1 condensates, exhibiting faster and more complete recovery of fluorescence after photobleaching (Figure 1D-E; Supplementary Figure 1D). Moreover, polyIC-induced HCoV-OC43/MERS-CoV N condensates displayed faster dynamics than polyIC-induced SARS-CoV/SARS-CoV-2 N condensates (Figure 1E, Supplementary Figure 1D). Overall, the seven HCoV N proteins exhibit varied basal phase separation propensities with constitutive condensates being less dynamic than polyIC-induced condensates, and in all cases, polyIC increased N condensation.

To gain insight into the specific regions of N that contribute to differences in basal phase separation behavior between species, we expressed ‘domain swap’ mutants of the SARS-CoV-2 and HCoV-229E N proteins in cells (Supplementary Figure 1E). We observed that individually replacing the SARS-CoV-2 N protein N-terminal domain (NTD) and central Ser/Arg (SR)-rich linker domains with the equivalent domains from HCoV-229E N results in a significant increase in basal phase separation propensity, suggesting that differences in the properties of the NTD and linker domains may explain the varied basal phase separation between the two N proteins.

### High-content phenotypic screening for modulators of SARS-CoV-2 N condensation

We hypothesized that modulating the phase behavior of SARS-CoV-2 N with small molecules may exert antiviral effects by perturbing the finely tuned dynamics of N condensates required for various stages of viral replication (Figure 2A). To identify compounds that promote or inhibit condensation of SARS-CoV-2 N, we devised two parallel screens (Figures 2B-C, Supplementary Table 1). Briefly, to identify compounds that promote N condensation (henceforth referred to as pro-condensers), A549 cells stably expressing SARS-CoV-2 N-EGFP were treated with 10μM compounds for 24h (Figure 2B). To identify condensate inhibitors, 17h after compound addition, cells were treated with polyIC for 7h (Figure 2C). After fixing and staining, cells were imaged at 20X magnification and the number of N puncta per cell was scored by image analysis (Supplementary Figure 2A). Positive control compounds were MS023, a type I protein arginine methyltransferase inhibitor known to induce N condensation (36), and salvianolic acid B (SalB), a natural product identified in our pilot screen that robustly inhibited formation of polyIC-induced condensates. The Z’ values for the pro-condensation and condensate inhibition screening modalities were 0.61 and 0.55 respectively. We performed both screening modalities against an annotated compound library comprising 2,082 FDA-compounds and 472 additional bioactive compounds at 10μM in technical duplicate, with two biological replicates. This was followed by confirmation of compounds in dose response experiments with both the original screening assay (Supplementary Figure 2A) and separate follow up experiments with an independent image analysis pipeline (Supplementary Figure 2B).

**FIGURE 2.**
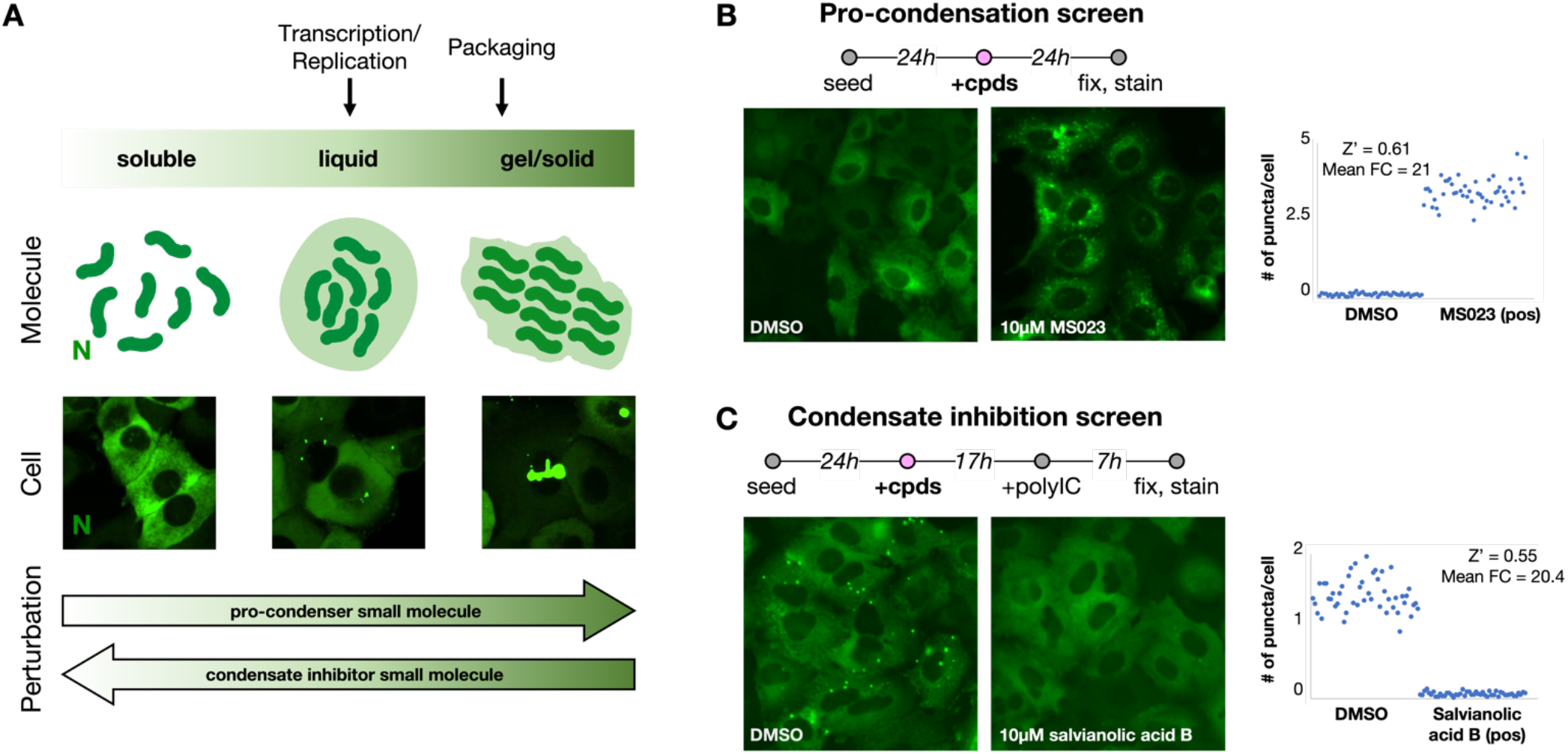
High content phenotypic screening assay to identify modulators of SARS-CoV-2 N condensation. (A) Schematic illustrating the sliding scale of N condensate material properties, adapted from Boeynaems et al., 2018 (1). N can exist in a soluble, diffuse cytoplasmic state, or anywhere along a continuum ranging from liquid-like condensates to more solid gel-like states. These various states are associated with different stages in viral replication such as transcription/replication and virion packaging. Intermolecular interaction strength tunes the material properties of condensates. Molecularly, N (as well as other components of N condensates) are more closely packed as interaction strength increases, leading to the formation and progression of liquid-like condensates to more gel-like aggregates. On a cellular level, liquid-like condensates appear as small spherical puncta, while progression to more gel-like states may result in the formation of larger, more irregularly-shaped aggregates. Small molecules that perturb the material state of N are referred to in this study as ‘pro-condensers’ and ‘condensate inhibitors’. (B) Schematic illustrating the pipeline for the pro-condensation screen to identify N pro-condensation compounds. Cells are seeded, then treated with compounds for 24h before fixing and staining. Representative images show the negative (DMSO) and positive (10μM MS023) controls. Quantification of number of N puncta per cell for controls is illustrated in the chart. Data indicates individual replicates. Images taken at 20x air magnification, widefield. Z’ for the screening assay is calculated as indicated in Methods. (C) Schematic illustrating the pipeline for the N condensate inhibition screen to identify N condensate-inhibiting compounds. Cells are seeded, treated with compounds for 17h, then treated with 1μg/mL transfected polyIC for an additional 7h before fixing and staining. Representative images show the negative (DMSO) and positive (10μM salvianolic acid B) controls. Quantification of number of N puncta per cell for controls is illustrated in the chart. Images taken at 20x air magnification, widefield. Z’ for the screening assay is calculated as indicated in Methods.

### SARS-CoV-2 N pro-condensation screen identifies GSK3 and proteasome inhibitors

After counter-screening to remove fluorescent artifacts, cytotoxic hits and compounds that act directly on EGFP, we identified six hit compounds that robustly increase the number of N puncta per cell (Figure 3A-B). These fell into two classes by annotation and follow-up: inhibitors of the proteasomal catalytic core complex and GSK3 inhibitors. Proteasome inhibition could prevent N turnover, thereby increasing the concentration of N in cells and promoting its phase separation. However, we also observed an increase in N nuclear localization upon treatment of cells with the proteasome inhibitors (Supplementary Figure 3A), suggesting that these small molecules may exert modulatory effects on N condensation through multiple mechanisms. Proteasome inhibition for cancer treatment has toxic side effects which preclude this target for anti-viral drugs, so this hit class was not pursued further.

**FIGURE 3.**
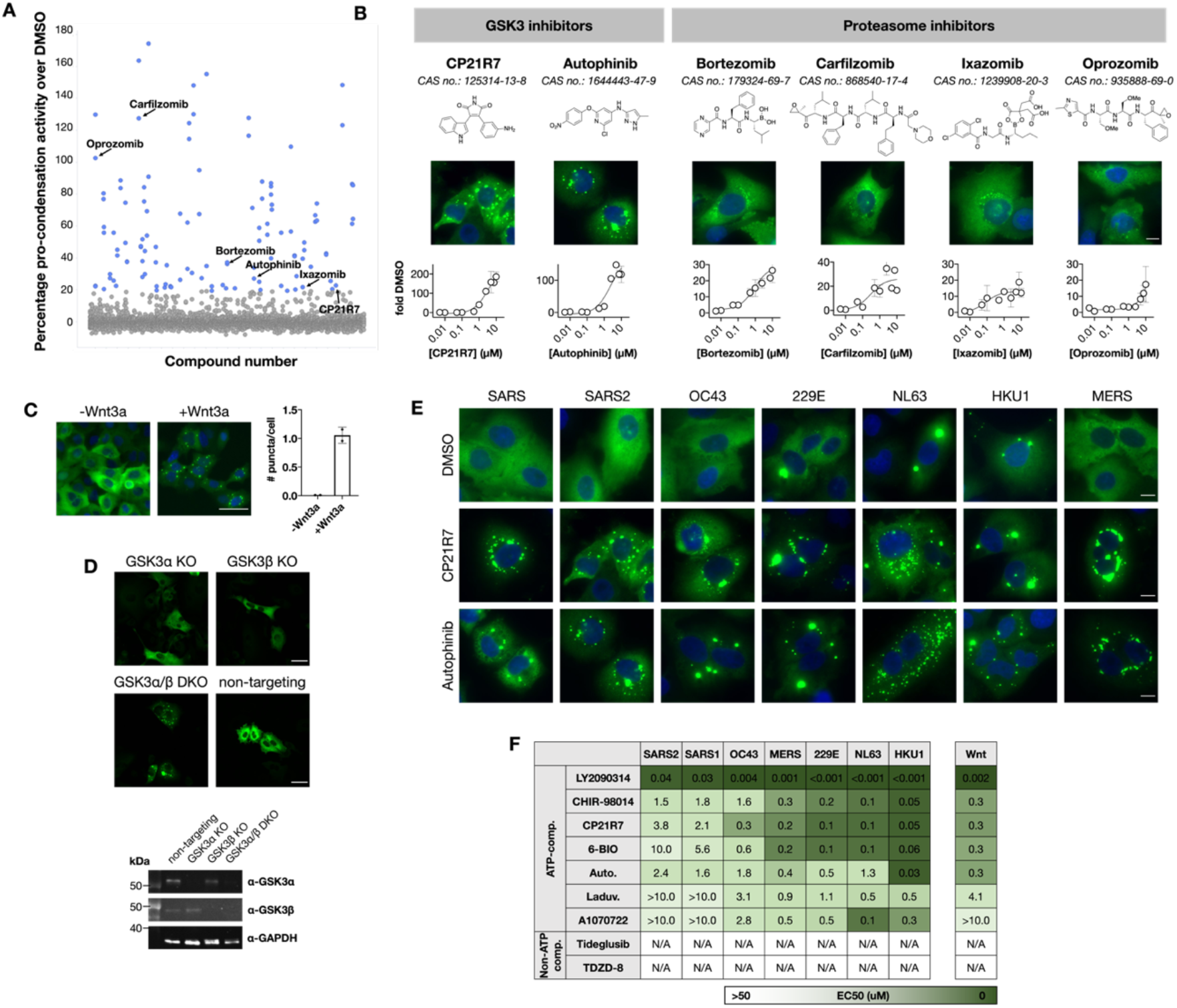
GSK3 and proteasome inhibitors promote pan-HCoV N condensation and condensate hardening. (A) Scatter plot illustrating SARS-CoV-2 N pro-condensation screening results. Data is represented as percentage N condensation activity. Data graphed shows both technical replicates. Blue points: identified compounds that showed N condensation activity greater than two standard deviations from the mean based on DMSO control percentage activity (see Methods for definition). Validated hit compounds are annotated on the plot. (B) Hit compounds classified according to their annotated cellular targets. Images are taken at 40x air magnification, widefield. Dose response curves indicate fold change in number of N puncta per cell over DMSO control (fold DMSO) and illustrate mean ± standard deviation of duplicates. Scale bar, 10 μm. (C) Fluorescence images and quantification for A549 SARS-CoV-2 N-EGFP cell lines treated with 400ng/μL Wnt3a for 24h. Graph illustrates mean ± standard deviation of duplicates. Images are taken at 20x air magnification, widefield. Scale bar, 20 μm. (D) Fluorescence images and western blot quantification for GSK3α, GSK3β and GSK3α/β CRISPR knockout cell lines illustrating increase in N condensation. Images are taken at 20x air magnification, widefield. Scale bar, 20 μm. Non-targeting: non-targeting sgRNA control. (E) Fluorescence images of all seven HCoV N-EGFP cell lines treated with 10μM GSK3 inhibitor hit compounds for 24h. Images are taken at 40x air magnification, widefield. Scale bar, 10 μm. (F) Left: N condensation EC50s for all seven HCoV N-EGFP cell lines across all seven ATP-competitive and two non-ATP-competitive GSK3 inhibitors. Right: EC50s for SARS-CoV-2/HCoV-NL63 N condensation and Wnt signaling as represented by β-catenin activation. β-catenin activation was measured using a MEF β-catenin-activated luciferase reporter cell line. EC50s shown are representative of at least two biological duplicate experiments. Auto., autophinib; Laduv., laduviglusib.

We identified one ATP-competitive GSK3 inhibitor (CP21R7) as a pro-condenser hit (Figure 3B). Autophinib, a second pro-condenser hit originally annotated as a VPS34 ATP-competitive inhibitor of autophagy, robustly induced N condensation and was later found to be a GSK3 inhibitor (see below). We therefore tested five other ATP-competitive GSK3 inhibitors (6-BIO, laduviglusib, A1070722, CHIR-98014, LY2090314) as well as the Mg^2+^-competitive GSK3 inhibitor lithium chloride (LiCl) and found them to also induce N condensation, albeit with varying EC50s that spanned several orders of magnitude, with the most potent compound being LY2090314 (Supplementary Figures 3B-C). Mass spectrometry analysis of N phosphorylation upon 1μM LY2090314 treatment confirmed inhibition of phosphorylation predominantly at the start of the SR-rich region of the LKR domain as well as at four other minor phosphorylation sites within the disordered N- and C-terminal arms and the N-terminal RNA-binding domain (Supplementary Figure 3D). Conversely, 1μM CP21R7 inhibited phosphorylation of N to a smaller extent, consistent with the different EC50s and N condensation potencies for these two small molecules.

To test if GSK3 is indeed the relevant target of the pro-condenser compounds, we pursued gain- and loss-of-function experiments. We observed robust SARS-CoV-2 N condensation when cells were stimulated with Wnt3a ligand, which activates the Wnt signaling pathway and leads to GSK3 inhibition (Figure 3C). CRISPR-knockouts of GSK3α/β recapitulated the effect of inhibitors, with CRISPR-knockouts of either kinase alone exhibiting a smaller effect than the double knockout (Figure 3D). Finally, site-directed mutagenesis of the 14 Ser residues within the SR-rich LKR region also recapitulated small molecule-induced condensation of N (Supplementary Figure 3E), further validating the on-target activity of these compounds. These Wnt pathway and genetic data confirm GSK3 as the target of the small molecule inhibitors and suggest that GSK3α and GSK3β play partially redundant roles in phosphorylating SARS-CoV-2 N and preventing its condensation, as they do for other proteins that are regulated by GSK3 (37).

### ATP-competitive GSK3 inhibitors induce pan-HCoV N condensate hardening

GSK3 was previously reported to regulate N from SARS-CoV and SARS-CoV-2 (27,29,38,39), but less closely related coronaviruses have not been tested. We thus followed up by treating all seven A549 cell lines expressing N-EGFP from the various human coronaviruses with all seven ATP-competitive GSK3 inhibitors and analyzing N condensation. Robust and reproducible dose dependent N condensation was observed across all seven Ns (Figure 3E-F, Supplementary Figure 4A-B). These data suggest that regulation of N condensation by GSK3 is conserved among HCoVs, even though the sequences of divergent Ns are only ~25% identical. We also found that treatment of N proteins from the bat coronaviruses (bat-CoVs) RaTG13, WIV1, HKU4, HKU10 and HKU8 with the most potent GSK3 inhibitor, LY2090134, induced their phase separation (Supplementary Figure 4C). Overall, our data indicates that N condensation across all seven HCoVs as well as several bat-CoVs is negatively modulated by GSK3. However, the sensitivity of different HCoV Ns to the pro-condensation effects of GSK3 inhibitors varied considerably (Figure 3F). For example, HCoV-HKU1 and HCoV-NL63 N were much more sensitive to compound modulation than SARS-CoV-2 N, with EC50s for all compounds typically being one to two orders of magnitude lower compared to SARS-CoV-2 N, despite similar N expression levels (Supplementary Figure 1A).

Dephosphorylation of SARS-CoV-2 N by inhibition of GSK3 is thought to promote N phase transition from a liquid-like condensate state to a more gel-like, less dynamic state (29). We probed the dynamics of various HCoV N condensates in the absence or presence of our most potent GSK3 inhibitor LY2090314, using FRAP. We observed that LY2090314-induced SARS-CoV, SARS-CoV-2, HCoV-OC43 and HCoV-229E N condensates were less dynamic than their corresponding polyIC-induced condensates, with percentage recovery over two minutes decreasing from 53.3%/50.5%/70.8%/72.1% in the polyIC-treated condition to 6.8%/13.9%/35.6%/5.6% in the LY2090314-treated condition, respectively (Supplementary Figure 4D). LY2090314 treatment did not result in any statistically significant changes in percentage recovery for constitutive HCoV-229E, HCoV-NL63 and HCoV-HKU1 condensates, likely owing to the already slow dynamics of constitutive N condensates. Overall, this suggests that ATP-competitive GSK3 inhibitors are not only capable of inducing N aggregation from a basal soluble state, but also result in hardened N condensates with much slower dynamics compared to polyIC-induced N condensates.

GSK3 has been considered as a therapeutic target for treatment of coronavirus infections (38–41), but a concern is possible toxicity. For GSK3-targeting inhibitors, the most relevant host pathway to consider for safety is canonical Wnt signaling through β-catenin, which is activated by GSK3 inhibition (42). This can drive hyperproliferation of epithelial cells in the gut, which is considered a negative safety signal (43). As a preliminary indicator of therapeutic index, we compared the inhibitor EC50 values obtained in our N condensation assays to EC50s for Wnt pathway activation. In addition to the ATP-competitive GSK3 inhibitors listed above, we also tested two non-ATP competitive inhibitors (tideglusib, TDZD-8), of which tideglusib has been shown to not activate β-catenin signaling (44). We found a correlation between the EC50s for N condensation and Wnt signaling activation (via β-catenin activation) for the ATP-competitive inhibitors, with more potent pro-condensation GSK3 inhibitors such as LY2090314 also activating Wnt signaling at lower concentrations (Figure 3F, right). Conversely, the non-ATP competitive inhibitors were inactive on both assays (Figure 3F; Supplementary Figure 4A). These data suggest that for HCoVs like SARS-CoV-2, it will be difficult to separate the safety risk of Wnt signaling activation from N modulation, since inhibitor concentrations required for N modulation would also activate Wnt signaling. However, therapeutic modulation of N condensation may be viable in the case of HCoVs whose Ns are unusually sensitive to GSK3 inhibition, such as HCoV-NL63 and HCoV-HKU1, where drug exposure below the threshold for activating Wnt signaling might be anti-viral.

### SARS-CoV-2 N condensate inhibitor screen identifies compounds that inhibit the polyIC input

Our condensation inhibition screen probed the same compound library. After counter-screening, we identified four hit compounds that robustly reduced the number of polyIC-induced N puncta per cell. These included three compounds annotated as Bcr-Abl/Src inhibitors (bosutinib, ponatinib and olverembatinib) as well as a plant-derived polyphenol, salvianolic acid B (SalB) (Figures 4A-B).

**FIGURE 4.**
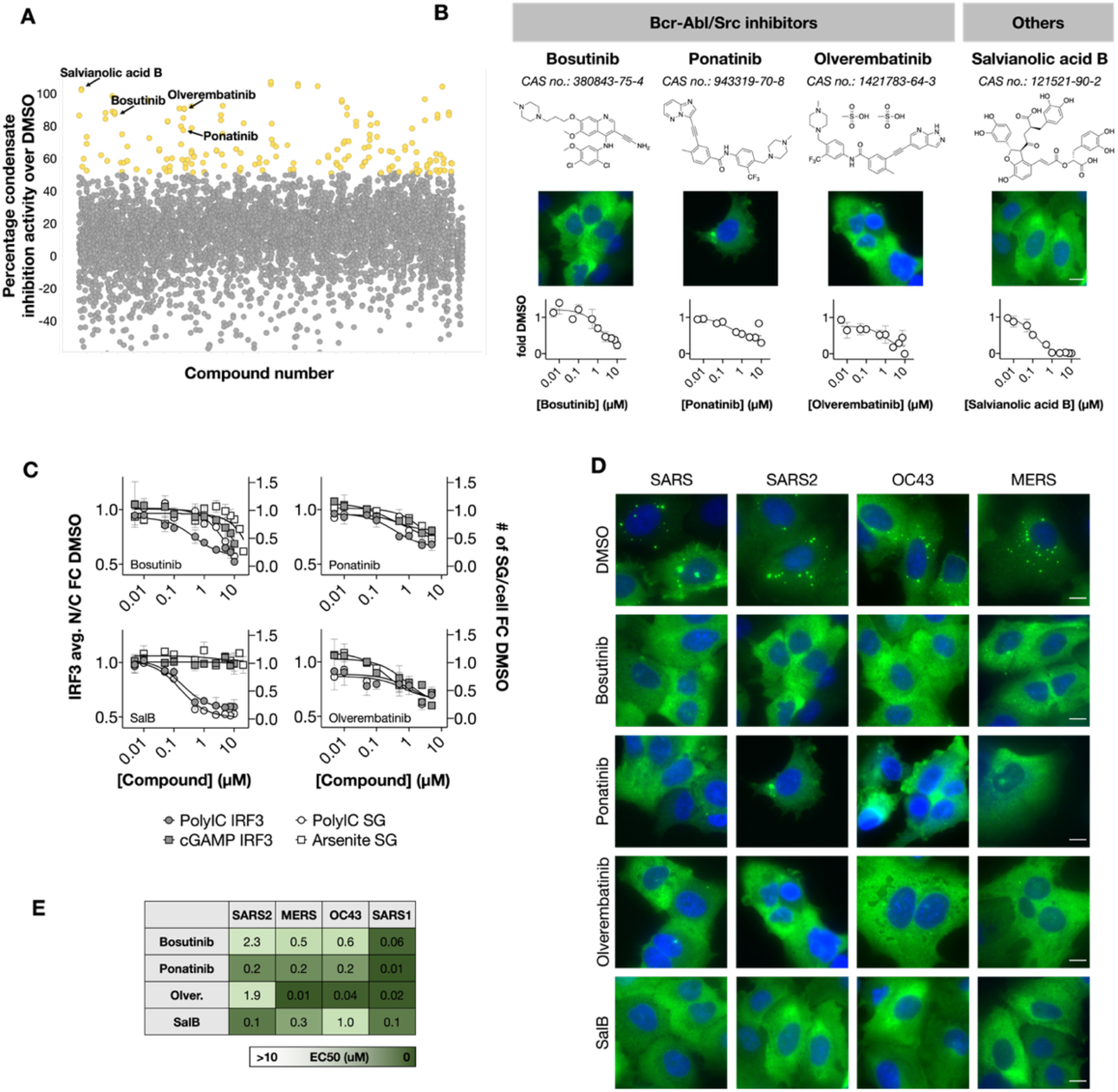
Diverse classes of compounds inhibit SARS-CoV-2 polyIC-induced N condensation. (A) Scatter plot illustrating SARS-CoV-2 N condensate inhibition screening results. Data is represented as percentage N condensate inhibition activity. Data graphed shows both technical replicates. Yellow points: identified compounds that showed N condensation inhibition activity greater than two standard deviations from the mean based on DMSO control percentage activity (see Methods for definition). Validated hit compounds are annotated on the plot. (B) Hit compounds classified according to their annotated cellular targets. Images are taken at 40x air magnification, widefield. Dose response curves indicate fold change of number of N puncta per cell over DMSO control (fold DMSO) and illustrate mean ± standard deviation of duplicates. Scale bar, 10 μm. (C) Dose response curves illustrating inhibition of stress granule formation and IRF3 activation by various N condensate inhibitors. Data shown indicates the fold change in average nuclear to cytoplasmic signal of IRF3 compared to DMSO (left y-axis) and the fold change in number of G3BP1-containing stress granules per cell compared to DMSO (right y-axis). All data indicate mean ± standard deviation of duplicates. (D) Representative fluorescence images of SARS-CoV/SARS-CoV-2/HCoV-OC43/MERS-CoV N-EGFP cell lines treated with 10μM condensate inhibitors 17h followed by 1μg/mL transfected polyIC treatment for an additional 7h. Images are taken at 40x air magnification, widefield. Scale bar, 10 μm. (E) N condensate inhibition IC50s for SARS-CoV/SARS-CoV-2/HCoV-OC43/MERS-CoV N-EGFP cell lines across all four active condensate inhibitors. IC50s shown are representative of at least two biological duplicate experiments.

To test if these compounds act directly on N itself, or indirectly via a host factor, we tested whether they inhibited two endogenous pathways that are induced by transfection of polyIC, in particular, stress granule (SG) formation and IRF3 translocation into the nucleus triggered by RIG-I and related viral RNA sensors. Both pathways were measured using cell-based high content assays. At high concentrations, the three annotated kinase inhibitors (bosutinib, ponatinib and olverembatinib) inhibited SG formation triggered by polyIC or arsenite, as well as IRF3 nuclear localization triggered by polyIC or cGAMP (Figure 4C, Supplementary Figure 5A). These data suggest action by polypharmacology. Bosutinib exhibited more potent activity against polyIC-triggered IRF3 translocation, suggesting a possibly interesting off-target activity on that pathway. We also tested two other Bcr-Abl kinase inhibitors (nilotinib, imatinib) and two other Src kinase inhibitors (saracatinib, PP2) in our assay. None of the compounds resulted in robust dose-dependent inhibition of polyIC-induced N condensation (Supplementary Figure 5B), suggesting that the relevant target(s) may not be Abl or Src family kinases. Interestingly, treatment with nilotinib instead resulted in an increase in N condensation, suggesting an additional possible off-target N condensation mechanism. The polyphenol SalB was more specific, inhibiting only polyIC-induced SG formation and IRF3 nuclear localization (Figure 4C, Supplementary Figure 5A). Thus, all the condensate inhibitors act against the polyIC input into the assay, presumably by inhibiting host factors required for polyIC signaling. We suspect the kinase inhibitors block the polyIC input via kinase inhibition, but likely not by inhibition of their annotated primary targets Abl or Src.

We next sought to determine the species-specificity of the four active condensate inhibitors (bosutinib, ponatinib, olverembatinib, SalB). All four compounds also showed significant suppression of polyIC-induced SARS-CoV/SARS-CoV-2/HCoV-OC43/MERS-CoV N condensation (Figure 4D-E, Supplementary Figure 5C). However, we did not observe significant inhibition of the formation of constitutive HCoV-229E/HCoV-NL63/HCoV-HKU1 N condensates (Supplementary Figure 5D), further demonstrating that the activity of the condensate inhibitors is polyIC-dependent. No conclusive inhibition of polyIC-induced N condensates could be determined for HCoV-229E/HCoV-NL63/HCoV-HKU1 owing to the relatively small difference in number of N puncta per cell between the polyIC-induced state and basal condensation state (Supplementary Figure 5E).

### Condensate formation and anti-viral activity

The library we screened contains approved drugs and well-annotated tool compounds. Similar libraries have been screened by multiple groups for antiviral activity against HCoVs (45–50). Several of the GSK3, Src/Abl and proteasome inhibitors we identified as N condensation modulators were previously shown to have antiviral activity. The published data cover several HCoV species and several different cell lines (Figure 5A). In addition, for several of the GSK3 inhibitors, published antiviral IC50s showed correlation with the N condensation EC50s determined here; notably, the most potent pro-condenser compound LY2090314 also exhibits antiviral activity against HCoV-229E at low inhibitor concentrations (Figure 5B). This suggests that possible on-target activity on N condensation may be responsible for the antiviral activity of the GSK3 inhibitors, and that small molecule modulation of N condensation can exert antiviral activity.

**FIGURE 5.**
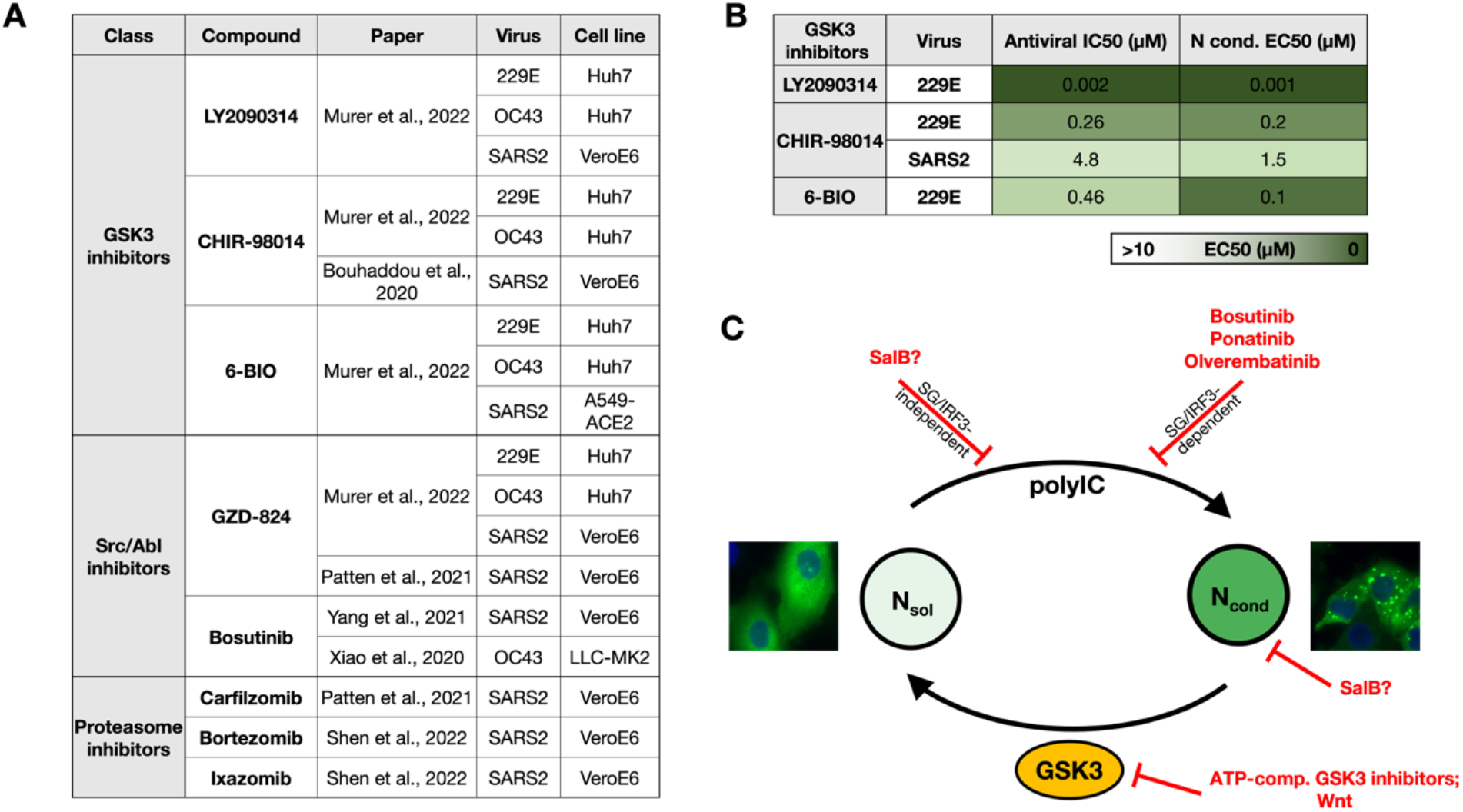
N condensate modulators are antiviral against SARS-CoV-2, HCoV-OC43 and HCoV-229E. (A) Active compounds from our screens and follow-up assays that have been identified as antiviral across various HCoV infection models by prior studies. (B) Antiviral IC50s (from prior studies, where available) and corresponding N condensation EC50s (from our N condensation assays) for GSK3 inhibitors illustrating a correlation between N condensation potency and antiviral activity. (C) Schematic illustrating the tunability of N condensation by small molecules identified in our screens. GSK3 phosphorylates all seven HCoV N proteins. ATP-competitive GSK3 inhibitors and Wnt signaling inhibit N phosphorylation and promote N condensation, with resultant condensates exhibiting significantly reduced dynamics. Inhibition of one or more host factors involved in SG/IRF3 signaling pathways by the Bcr-Abl/Src kinase family of inhibitors prevent polyIC/RNA-induced N condensation. In addition, SalB may inhibit polyIC/RNA-induced N condensation through different mechanisms such as the inhibition of host factor(s) that induce N condensation through SG/IRF3-independent mechanisms, and/or direct interference of N-RNA/polyIC interactions.

## Discussion

Biomolecular condensates play a role in cellular processes such as embryonic development, stress response and pathological aggregation of proteins, and are also critical for various stages of viral replication. In this study, we show that (1) N condensation is a common phenomenon across all seven HCoVs, and (2) small molecules can promote or inhibit N condensation via perturbation of host targets, and this activity tends to be common across N proteins from all HCoV species tested (Figure 5C). Several of these small molecules are also active against multiple HCoV infection models. These data show that perturbation of viral condensate dynamics via host factors has the potential to generate drugs with antiviral activity across multiple viral species, including new pandemic species. Our approach also illustrates that cell-based screens using viral genes can predict potential anti-viral activity of small molecules without requiring access to whole virus infection models.

Our high content screens identified small molecule inhibitors of GSK3 that tune both the fraction of condensed N protein as well as the dynamics of N condensates. GSK3 has previously been proposed as a HCoV target (39–41). Our work reveals its pan-HCoV potential, but also highlights the safety risk associated with Wnt pathway activation. GSK3 is an abundant, constitutively active Ser/Thr kinase that phosphorylates a wide range of pre-primed substrates (51) and has previously been shown to phosphorylate both SARS-CoV and SARS-CoV-2 N along its SR-rich LKR region (27,29,38). In this study, we show that GSK3 inhibitors exhibit the same condensate modulating effects across all seven HCoV and five bat-CoV N proteins. These condensate-hardening inhibitors likely inhibit viral replication through on-target induction of N condensation/aggregation. In addition, several others have also demonstrated that GSK3 inhibitors are antiviral against various HCoVs (38,39,45,47). This further promotes GSK3 as candidate target for development of multi-CoV anti-virals and illustrates one of the major benefits of host-targeting to achieve broad-spectrum anti-viral activity (in addition to reduced risk of resistance development). However, as with any host target, a key concern is therapeutic index and toxicity due to perturbation of host pathways that depend on the target, in this case potentially oncogenic β-catenin signaling. For the HCoVs whose Ns are much more sensitive to compound modulation such as HCoV-NL63, it could be that anti-viral activity can be achieved with minimal activation of β-catenin signaling in live virus infection models. In addition, a follow-up would be to investigate the molecular basis behind the difference in sensitivity of various Ns to compound modulation, for example by determining if dephosphorylation occurs at lower concentrations of GSK3 inhibitors for more sensitive N proteins, or if these N proteins dephosphorylate at similar concentrations of inhibitor but require a lower degree of dephosphorylation to condense. Condensate targets offer novel avenues for optimization chemistry, notably the potential to increase on-target activity by partitioning of the drug into the condensate (52). Since viral and host condensates have different compositions, this effect should enable improvement of selectivity. We compared EC50 values for nine GSK3 inhibitors in N condensate versus Wnt activation assays and found no compounds that were notably selective for the viral pathway over the host pathway (Figure 3F). Testing a larger library of GSK inhibitors, or a focused medicinal chemistry effort, might tease out selectivity between these pathways.

The N condensate inhibitors we identified appear to target the polyIC input to N condensation, as evidenced by their ability to block induction of polyIC-induced IRF3 translocation and SG assembly. A potentially causal relationship between SG induction and SARS-CoV-2 N condensation has been shown by others (21,53). The three kinase inhibitors we identified (bosutinib, ponatinib and olverembatinib) are annotated as targeting Bcr-Abl and Src. However, our investigation of additional potent Bcr-Abl and Src inhibitors failed to support this hypothesis. We currently suspect their activities, especially at concentrations of 1μM and higher, may be due to polypharmacology that could be resolved by kinase activity profiling and additional SAR.

The polyphenolic natural product SalB represents a class of compounds that has a broad range of biological effects and are considered not promising starts for medicinal chemistry. SalB selectively blocks polyIC input at some stage as evidenced by its inhibition of only polyIC-induced IRF3 activation and SG formation. However, its polypharmacology raises the possibility that it may also act directly on N by physically inhibiting N-RNA or general RNA-protein interactions. It was shown in a previous study that another polyphenol natural product, (-)-gallocatechin gallate, is able to disrupt SARS-CoV-2 N condensation through direct interference of N-RNA binding (54). Taken together, the condensate inhibitors demonstrate that small molecules are capable of disrupting liquid-liquid phase separation of HCoV N proteins through varied mechanisms.

The use of the FDA approved library that has been screened several times for anti-viral activity allows us to see a potential correlation with condensate formation and virus infectivity. However, one of the limitations of screening with an annotated FDA-approved library is the lack of diversity in pharmacological targets. Screening a larger, more chemically diverse compound library is a natural next step for our approach. Additionally, while our primary goal in screening for modulators of HCoV N condensates was to identify targets for treating COVID-19 and other HCoV infections, our approach is also relevant to other diseases where ribonucleoprotein (RNP) aggregates have been causally implicated. Neurological diseases such as Huntington’s disease, spinocerebellar ataxia and Fragile X syndrome arise from nucleotide repeat expansions in non-coding RNA that give rise to pathological nuclear RNP granules (55). Similarly, pathological cytoplasmic RNP inclusions of mutant variants of the Fused in Sarcoma (FUS) protein are the hallmark of ALS (56–58). In addition to its role in virus infections, double-stranded RNA signaling may also underly neurodegeneration caused by the *C9ORF72* locus (59). Taken together, the ability to screen for condensate dynamics as a therapeutic target supports the value of moving forward with larger more diverse compound libraries to reveal novel condensate biology in both viral infections as well as in other indications such as neurological diseases.

## Materials and Methods

### Cell lines and cell culture

HEK293T/17 (CRL-11268) and A549 (CCL-185) cells were purchased from ATCC. BJ-5ta ΔcGAS cells were obtained from Dr. Tai L. Ng (Harvard Medical School). HEK293T/17 cells and BJ-5ta ΔcGAS cells were maintained in Dulbecco’s Modified Eagle Medium (DMEM; ATCC 30-2002) supplemented with 10% fetal bovine serum (FBS; Gibco 10438026), 1,000U/mL penicillin-streptomycin (Gibco 15140122) and 100μg/mL normocin (Invivogen ant-nr-1). Wild-type A549 cells were maintained in F-12K medium (ATCC 30-2004) supplemented with 10% FBS, 1,000U/mL penicillin-streptomycin and 100ug/mL normocin or DMEM supplemented with 10% FBS. A549 stable cell lines expressing various HCoV N-EGFP were maintained in full F-12K culture medium with 1.5μg/mL puromycin (Gibco A1113803). Cells were maintained at 37°C and 5% CO_2_ in a humidified environment and subcultured twice a week by DPBS washing (Gibco 14190250) followed by trypsinization (Corning MT25053CI) from 90% to 20% confluence.

### Plasmid construct generation

The pHAGE lentiviral plasmid encoding SARS-CoV-2 N-EGFP was obtained from Dr. Adrian Salic (Harvard Medical School). Plasmid vectors containing SARS-CoV, HCoV-OC43, HCoV-229E, HCoV-NL63, HCoV-HKU1 and MERS-CoV N sequences were obtained from Dr. Tai L. Ng (Harvard Medical School). pHAGE lentiviral plasmids encoding the six other HCoV N-EGFP were generated by replacing SARS-CoV-2 N with the respective HCoV N sequences by PCR (New England Biolabs M0492S) and NEBuilder HiFi DNA Assembly (New England Biolabs E2621S). Oligonucleotides used for PCR were obtained from Genewiz, and plasmids were verified by Sanger sequencing at Genewiz. The pHAGE plasmid encoding SARS-CoV-2 N^SAmut^-EGFP was synthesized by Twist.

### Stable A549 cell line generation

Stable A549 cell lines expressing the seven HCoV N-EGFP were generated by lentiviral transduction followed by 1.5ug/mL puromycin selection. 12μg of each pHAGE lentiviral construct encoding the various HCoV N-EGFP were co-transfected with 18μg of the psPAX2 (Addgene #12260) and 6μg of the pMD2.G (Addgene #12259) 2^nd^ generation lentiviral packaging plasmids into HEK293T/17 cells at 70% confluency in T75 flasks with Lipofectamine3000 (Invitrogen L3000015). After 6-24h, transfection medium was removed and replaced with full DMEM. Culture medium containing lentivirus was collected after a subsequent 24h and 48h and combined. Virus-containing medium was centrifuged at 400xg for 5 min to pellet cell debris before filtering through a 0.45μm Durapore PVDF membrane Steriflip (Millipore SE1M003M00). 4X Lenti-X Concentrator (TaKaRa 631232) was then added to the filtered virus-containing medium, incubated at 4°C for 1h, then centrifuged at 1,500xg for 45 min. After centrifugation, the supernatant was removed and the virus pellet was resuspended in 1/10^th^ the original volume of harvested supernatant, in complete medium without antibiotics. Two days before transduction, 250,000 A549 cells were plated in each well of a 6-well plate, and transductions were performed by replacing wells with 1.5mL of concentrated virus, 0.4mL of 10X polybrene (Millipore TR-1003-G) and 2.1mL of complete medium without antibiotics. Cells were incubated for 36-48h before virus removal and expansion into 1.5μg/mL puromycin-containing selection medium. After selection recovery, stable pools of cells were analyzed by qualitative fluorescence microscopy and expanded for liquid nitrogen storage in 90% FBS, 10% DMSO.

### SDS-PAGE and western blots

Cells were seeded at 10,000 cells/well in 96-well plates and treated accordingly. Cells were washed once with ice-cold PBS, then lysed in 50μL ice-cold lysis buffer comprising 25mM Tris-HCl pH7.4, 150mM NaCl, 1% Triton X-100 (Millipore X100), 1X Halt protease inhibitor (Thermo Scientific 87785) and 1U/μL Benzonase (Millipore E1014). Samples were boiled with an equal volume of 2X Laemmli sample buffer (Bio-Rad 1610737) containing β-mercaptoethanol for 10 min before running on an SDS-PAGE gel (Invitrogen XP10205BOX). Gels were transferred onto nitrocellulose membranes using the iBlot 2 Dry Blotting System (Invitrogen IB23002) and blocked in SuperBlock™ (TBS) Blocking Buffer (Thermo Scientific 37535) for 1h at RTP. Primary antibodies (HRP anti-GFP antibody [Abcam ab190584]; rabbit anti-GSK3α antibody [Cell Signaling Technology 4818S]; rabbit anti-GSK3β antibody [Cell Signaling Technology 9315S]) were diluted in Pierce™ Protein-Free T20 (TBS) Blocking Buffer (Thermo Scientific 37571) as recommended and incubated overnight at 4°C, shaking. Membranes were then washed thrice with TBS + 0.2% Tween-20. For HRP anti-GFP antibody samples, membranes were treated with SuperSignal™ West Femto Maximum Sensitivity Substrate (Thermo Scientific 34095) before visualization with a Bio-Rad imager. For all other antibodies, membranes were incubated with goat anti-Rabbit IgG (H+L) Secondary Antibody, DyLight™ 680 (Invitrogen 35568) diluted in Pierce™ Protein-Free T20 (TBS) Blocking Buffer (Thermo Scientific 37571) as recommended for 1h at RTP. Membranes were then washed again thrice with TBS + 0.2% Tween-20 before visualization with a Bio-Rad imager.

### Fluorescence microscopy

For qualitative confocal fluorescence imaging, cells were imaged with a Nikon Ti fluorescence microscope equipped with a Yokogawa CSU-W1 spinning disk confocal scanner, Nikon LUN-F XL solid state laser combiner, Hamamatsu ORCA-Fusion BT CMOS camera, motorized stage and shutters and Lumencor SOLA fluorescence light source. Cells were seeded in 24-well or 96-well high performance #1.5 cover glass bottom plates (Cellvis P24-1.5H-N; P96-1.5H-N) at 100,000 or 10,000 cells/well in full medium without antibiotics and were typically imaged 24-48h after seeding. N-EGFP was imaged with a 488nm laser and ET525/50m filter (Chroma), while Hoechst staining was imaged with a 405nm laser and ET455/50m filter (Chroma). For qualitative widefield fluorescence imaging, cells were imaged with a Nikon Ti2 fluorescence microscope equipped with a Hamamatsu Flash 4.0 LT camera and a SOLA fluorescence light source. Cells were seeded in 96-well black wall, clear bottom plates (Corning 9603) at 10,000 cells/well in full medium without antibiotics and were typically imaged 24-48h after seeding. N-EGFP was imaged with a 466/40 excitation filter and a 525/50 emission filter.

### Fluorescence recovery after photobleaching (FRAP)

FRAP experiments were performed on a DeltaVision OMX Blaze microscope equipped with a 60x/1.42 Plan Apo oil objective (Olympus), a 488nm laser and a PCO edge Front Illuminated sCMOS camera. N-EGFP was imaged with an LED, 477/32nm bandpass filter and 528/48nm emission filter. Bleaching of N-EGFP was performed with a 488nm laser. For each experimental sample, seven condensates were bleached within a circular region of about 1.2 μm diameter at 31.3% laser transmission for 50 ms. A total of 120 frames were recorded at one frame per 2s, for a total of 2 min (two frames recorded prior to bleach event, followed by 117 subsequent post-bleach frames). Images were processed in FIJI software (NIH). The fluorescence intensity of a 0.32 μm-diameter circular region of interest (ROI) within the bleached spot was monitored over time (I_bleached_). The fluorescence intensity within the same ROI pre-bleaching (I_pre-bleach_) and immediately post-bleaching (I_post-bleach_) were also recorded. The fluorescence intensity of a separate rectangular ROI of 2.72 μm diameter away from one bleached ROI per sample was monitored over time (I_background_). The FRAP recovery intensities at any given time point (I_t_) were calculated as follows:

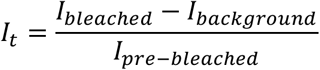

The curves obtained were then normalized as follows:

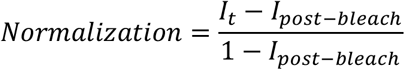

The mean and standard deviation of the final normalized values were plotted for seven condensates per experimental condition, and the final normalized recovery value was taken as the percentage recovery for each condition.

### High content compound screening and follow-up experiments

The primary screen was performed at AbbVie as follows. Briefly, cells were seeded in Perkin Elmer LLC ViewPlate-384 well black, optically clear bottom plates at 2,000 cells/well and incubated overnight. The Selleck compound library (2,554 compounds) was added at a final concentration of 10μM and incubated overnight on duplicate plates. Plates for the condensate inhibition screen were additionally transfected with a final concentration of 1ug/mL polyIC 17h after compound addition to induce N condensation and incubated for a further 7 hours. All plates for both the pro-condensation and condensation inhibition screens were fixed with 3% formaldehyde and nuclei stained with Hoechst 33342 for nuclear identification. Plates were scanned on the Thermo Fisher CX7 LZR using a 20x objective and widefield imaging mode. Nuclei staining was imaged with the 405LZR_BGFR_BGFR filter. GFP was imaged with the 488LZR_BGFR_BGFR filter. Images were analyzed with automatic image analysis as described below.

The Z’ for both the pro-condensation and condensation inhibition assays were calculated as follows:

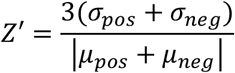

where σ_pos_ = SD of positive control, σ_neg_ = SD of negative control (DMSO), μ_pos_ = mean of positive control, μ_neg_ = mean of negative control (DMSO). For visualization of screening data (Figures 3A and 4A), % activity compared to DMSO control was calculated as follows:

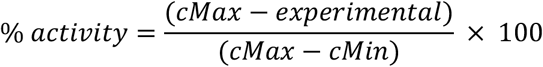

For the pro-condensation screen, cMax = mean number of puncta per cell for positive control MS023 and cMin = mean number of puncta per cell for negative control DMSO. For the condensation inhibition screen, cMax = mean number of puncta per cell for negative control DMSO and cMin = mean number of puncta per cell for positive control salvianolic acid B.

Follow-up experiments with hit compounds across all seven HCoV N-expressing A549 cell lines were performed at Harvard Medical School as above with slight modifications. All hit compounds were re-purchased for testing (see compound table below). Cells were seeded in 384-well PerkinElmer PhenoPlate 384-well black plates (PerkinElmer; 6057302) at 2,500 cells/well with the Thermo Multidrop™ Combi reagent dispenser and drugs were added in dose-response format with the HP D300e digital dispenser. After fixation and staining with 1μg/mL Hoechst 33342 (Thermo Scientific 62249) and 5μg/mL CellMask™ Deep Red Plasma membrane Stain (Invitrogen C10046), plates were were imaged with MetaXpress (version 6.7) on a Molecular Devices ImageXpress Micro Confocal Laser (IXM-C LZR) microscope equipped with a spinning disk confocal scanner, a high-quantum efficiency sCMOS detection camera and a Lumencor Celesta light engine. Cells were seeded in 96-well high performance #1.5 cover glass bottom plates (Cellvis P96-1.5H-N) or 384-well PerkinElmer PhenoPlate 384-well black plates (PerkinElmer 6057302) at 10,000 or 2,500 cells/well in full medium without antibiotics. Specimens were sampled with a 40x/0.95 air objective lens (Nikon) at a pixel size of 0.16μm/pixel. N-EGFP was imaged with a 477nm laser, 488nm dichroic filter and 520/25nm emission filter, while samples stained for immunofluorescence were imaged with a 546nm laser, 593nm dichroic filter and 624/40nm emission filter. Hoechst staining was imaged with a 405nm laser, 421nm dichroic filter and 452/45nm emission filter, while CellMask™ Deep Red Plasma membrane staining was imaged with a 638nm laser, 656nm dichroic filter and 692/40nm emission filter. All images were processed in FIJI software (NIH) and analyzed with automatic image analysis as described below.

### Automatic image analysis

Image analysis for the primary screen as well as follow-up experiments were performed at AbbVie and HMS respectively with independent image analysis pipelines that demonstrated highly reproducible results. At AbbVie, image analysis was performed using SpotDetector.V4 algorithm and is summarized in Supplementary Figure 2A. The output feature SpotCountPerObject was used to evaluate the changes in N protein aggregation in both screens. Compounds that showed activity greater than two standard deviations from the mean based on control % activity were selected. After visual inspection to remove artifacts and false positives, 25 pro-condensation and 44 condensate inhibiting compounds were tested using six-point CRC using the same assays described above. Six pro-condensers and four condensate inhibitors confirmed with EC50/IC50 and were identified as hits. At Harvard Medical School, image analysis for follow-up experiments was performed with Molecular Devices MetaXpress software with customized image analysis pipelines for quantification of N puncta and is summarized in Supplementary Figure 2B. For quantification of N puncta upon treatment with the pro-condensation proteasome inhibitors, thresholds for the image analysis pipeline were modified slightly to enable more robust and accurate identification of smaller and dimmer puncta. For dose response experiments, mean values were fit with a three-parameter curve in Prism (GraphPad 9).

### Mass spectrometry

For detection of N phosphorylation states, A549 SARS-CoV-2 N-EGFP cells were seeded in T-25 flasks at 40% confluency overnight before treatment with DMSO, 1μM LY2090314 or 1μM CP21R7 for a further 24h. Cells were then lysed with ice cold RIPA lysis buffer (supplemented with NaCl to a final concentration of 300mM) + 1X Halt protease and phosphatase inhibitor for 10min at 4°C. Cell lysates were then subjected to immunoprecipitation with GFP Selector resin (NanoTag Biotechnologies N0310-L) according to manufacturer’s protocol. After the final wash, the resin containing sample was resuspended in 50μL wash buffer before addition of 50μL 2X Laemmli sample buffer and subjected to SDS-PAGE. Gel bands were excised and submitted for mass spectrometry analysis at the Taplin Mass Spectrometry facility in Harvard Medical School (60). Data presented indicates percentage of each peptide that is phosphorylated as determined by peak intensity values.

### Generation of A549 SARS-CoV-2 N-EGFP GSK3α, GSK3β and GSK3α/β CRISPR knockout cells

Stable A549 SARS-CoV-2 N-EGFP GSK3α, GSK3β and GSK3α/β knockout cell lines were generated by lentiviral transduction of lentiCas9-Blast and GSK3α and/or GSK3β gRNAs followed by 1.5ug/mL puromycin and 10μg/mL blasticidin selection. lentiCas9-Blast was a gift from Feng Zhang (Addgene plasmid #52962) (61), while all GSK3α, GSK3β and non-targeting control gRNAs were a gift from John Doench & David Root (Addgene plasmids #77281, #77282, #77283, #76370, #76371, #76372, #80248, #80263) (62). Lentiviral pools containing GSK3α, GSK3β and non-targeting control gRNAs were generated as described earlier and transduced into the A549 SARS-CoV-2 N-EGFP cell line. After selection recovery, stable pools of cells were analyzed by quantitative fluorescence microscopy.

### Wnt pathway reporter assay

Mouse embryonic fibroblasts (MEFs) cells stably expressing a firefly luciferase gene under the control of TCF/LEF response element and Renilla luciferase under the control of a constitutive promoter were used to measure Wnt pathway activation (63). Cells were plated in 96-well plates and grown to confluence for 24h, and then treated in triplicate with the indicated GSK3 inhibitors in serum-free DMEM for 24 h. Luminescence was measured from cell lysates using the Dual-Glo Luciferase Assay System (Promega E1910) in a Victor3 Multilabel Plate Reader (Perkin-Elmer). Relative luciferase is represented as mean Firefly/Renilla ratio, with error bars representing standard deviation. Experiments were performed in dose response format, and mean values were fit with a four-parameter curve in Prism (GraphPad 9) to obtain EC50 values.

### Immunofluorescence staining for quantification of IRF3 nuclear/cytoplasmic ratios and SG formation

Cells were seeded in 384-well PerkinElmer PhenoPlate 384-well black plates (PerkinElmer 6057302) at 2,500 cells/well. To observe the effect of compounds on IRF3 activation and nuclear translocation, A549/BJ-5ta ΔcGAS cells were treated with compounds for a total of 17h/22h before addition of 1μg/mL transfected polyIC/100μg/mL cGAMP for an additional 7h/2h respectively. To observe the effect of compounds on stress granule formation, A549 cells were treated with compounds for a total of 17h/23h before addition of 1μg/mL transfected polyIC/0.5mM NaAsO2 for an additional 7h/1h respectively. Cells were then fixed with 4% paraformaldehyde for 20 min, washed thrice with 1X PBS and blocked in PBS + 0.1% Triton X-100 (Sigma-Aldrich X100) + 5% bovine serum albumin for 1h at RTP. Plates were then incubated overnight at 4°C with primary antibodies (rabbit anti-IRF3 antibody [Cell Signaling Technology 11904S]; rabbit anti-G3BP1 antibody [Abcam ab181149]) diluted in PBS + 0.1% Triton X-100 (Sigma-Aldrich X100) + 1% bovine serum albumin at recommended concentrations. Plates were washed thrice with 1X PBS before incubation with goat anti-Mouse IgG (H+L) Cross-Adsorbed Secondary Antibody, Alexa Fluor™ 568 (Invitrogen A-11004), 1μg/mL Hoechst 33342 and 5μg/mL CellMask™ Deep Red Plasma membrane Stain in the same dilution buffer for 1h at RTP. Plates were washed thrice again with 1X PBS before imaging with the IXM-C LZR as described earlier. Image analysis was performed with Molecular Devices MetaXpress software with customized image analysis pipelines for quantification of nuclear/cytoplasmic ratios of IRF3 and quantification of G3BP1 puncta. Briefly, for quantification of nuclear/cytoplasmic ratios of IRF3, nuclei and cytoplasm areas were defined by Hoechst and CellMask staining respectively, and the ratio of the average fluorescence intensity of IRF3 in the nucleus to cytoplasm was determined (total integrated intensity divided by total area) for each cell. For quantification of G3BP1 puncta, a customized pipeline similar to that utilized for N puncta quantification was used. Data for ponatinib and olverembatinib is presented up to 5μM concentration due to slight compound toxicity affecting the quality of image analysis.

### Compounds/stimuli for follow up experiments

All hit compounds were re-purchased for follow up experiments and validation at Harvard Medical School. Sources of all compounds used in this paper are shown in the table below. Sources of LiCl and Wnt3a ligand used in this paper are also indicated below.

| Compound/stimuli | Source | Catalog no. | Format |
| --- | --- | --- | --- |
| CP21R7 | MedChemExpress | HY-100207 | 10mM in 1mL DMSO |
| Autophinib | MedChemExpress | HY-101920 | 10mM in 1mL DMSO |
| Bortezomib | MedChemExpress | HY-10227 | 10mM in 1mL DMSO |
| Carfilzomib | MedChemExpress | HY-10455 | 10mM in 1mL DMSO |
| Ixazomib citrate | MedChemExpress | HY-10452 | 10mM in 1mL DMSO |
| Oprozomib | MedChemExpress | HY-12113 | 10mM in 1mL DMSO |
| 6-BIO | MedChemExpress | HY-10580 | 10mM in 1mL DMSO |
| Laduviglusib | MedChemExpress | HY-10182 | 10mM in 1mL DMSO |
| A1070722 | MedChemExpress | HY-107531 | 10mM in 1mL DMSO |
| CHIR-98014 | MedChemExpress | HY-13076 | 10mM in 1mL DMSO |
| LY2090314 | MedChemExpress | HY-16294 | 10mM in 1mL DMSO |
| Tideglusib | MedChemExpress | HY-14872 | 10mM in 1mL DMSO |
| TDZD-8 | MedChemExpress | HY-11012 | 10mM in 1mL DMSO |
| Bosutinib | MedChemExpress | HY-10158 | 10mM in 1mL DMSO |
| Olverembatinib dimesylate | MedChemExpress | HY-15666A | 10mM in 1mL DMSO |
| Ponatinib | MedChemExpress | HY-12047 | 10mM in 1mL DMSO |
| Salvianolic acid B | MedChemExpress | HY-N1362 | 5mg |
| Imatinib | MedChemExpress | HY-15463 | 10mM in 1mL DMSO |
| Saracatinib | MedChemExpress | HY-10234 | 10mM in 1mL DMSO |
| PP2 | MedChemExpress | HY-13805 | 10mM in 1mL DMSO |
| LiCl | Sigma-Aldrich | L9650 | 100g |
| Recombinant mouse Wnt3a (carrier-free) | R&D Systems | 1324-WN-010/CF | 10µg |

## Acknowledgements

This work was sponsored by a research alliance with AbbVie Inc. Rui Tong Quek was supported by the Agency for Science, Technology and Research NSS (PhD) predoctoral fellowship. The content and conclusions included in this document are solely those of the authors and do not necessarily represent official views of any funders. We thank the Nikon Imaging Center at HMS for help with fluorescence microscopy, the Taplin Mass Spectrometry Facility at HMS for help with mass spectrometry, and the ICCB-Longwood Screening Facility at HMS, in particular Dr. Clarence Yapp for assistance with image analysis and for access to high-throughput screening equipment.

R.T.Q. and K.S.H. generated cell lines and designed the high-throughput screening assay. S.G.W. performed the high-throughput small molecule screens and validation of hits. D.T.N. designed custom image analysis pipelines and assisted with image analysis. T.A.M. performed the Wnt reporter assays and processed the data. R.T.Q. performed qualitative imaging, follow up/mechanistic assays, data analyses and wrote the manuscript. S.M.G., P.A.S. and T.J.M. managed the team, assisted with data analysis and interpretation and provided scientific insight and advice on writing the paper. AbbVie participated in the study design and execution of experiments including the interpretation of data, review and approval of the publication. AbbVie provided financial support for this research. All authors provided critical feedback for the manuscript.

The authors declare no competing interests.

## References

1. Boeynaems, S., Alberti, S., Fawzi, N. L., Mittag, T., Polymenidou, M., Rousseau, F., Schymkowitz, J., Shorter, J., Wolozin, B., Van Den Bosch, L., Tompa, P., and Fuxreiter, M. (2018) Protein Phase Separation: A New Phase in Cell Biology. Trends Cell Biol 28, 420–435

2. Banani, S. F., Lee, H. O., Hyman, A. A., and Rosen, M. K. (2017) Biomolecular condensates: organizers of cellular biochemistry. Nature Reviews Molecular Cell Biology 18, 285–298

3. Shin, Y., and Brangwynne, C. P. (2017) Liquid phase condensation in cell physiology and disease. Science 357, eaaf4382

4. Zhao, Y. G., and Zhang, H. (2020) Phase Separation in Membrane Biology: The Interplay between Membrane-Bound Organelles and Membraneless Condensates. Developmental Cell 55, 30–44

5. Lahaye, X., Vidy, A., Pomier, C., Obiang, L., Harper, F., Gaudin, Y., and Blondel, D. (2009) Functional Characterization of Negri Bodies (NBs) in Rabies Virus-Infected Cells: Evidence that NBs Are Sites of Viral Transcription and Replication. J Virol 83, 7948–7958

6. Nikolic, J., Le Bars, R., Lama, Z., Scrima, N., Lagaudrière-Gesbert, C., Gaudin, Y., and Blondel, D. (2017) Negri bodies are viral factories with properties of liquid organelles. Nature Communications 8, 58

7. Heinrich Bianca, S., Maliga, Z., Stein David, A., Hyman Anthony, A., Whelan Sean, P. J., and Palese, P. Phase Transitions Drive the Formation of Vesicular Stomatitis Virus Replication Compartments. mBio 9, e02290–02217

8. Heinrich, B. S., Cureton, D. K., Rahmeh, A. A., and Whelan, S. P. J. (2010) Protein expression redirects vesicular stomatitis virus RNA synthesis to cytoplasmic inclusions. PLoS Pathog 6, e1000958–e1000958

9. Rincheval, V., Lelek, M., Gault, E., Bouillier, C., Sitterlin, D., Blouquit-Laye, S., Galloux, M., Zimmer, C., Eleouet, J.-F., and Rameix-Welti, M.-A. (2017) Functional organization of cytoplasmic inclusion bodies in cells infected by respiratory syncytial virus. Nature Communications 8, 563

10. Hoenen, T., Shabman, R. S., Groseth, A., Herwig, A., Weber, M., Schudt, G., Dolnik, O., Basler, C. F., Becker, S., and Feldmann, H. (2012) Inclusion bodies are a site of ebolavirus replication. J Virol 86, 11779–11788

11. Guseva, S., Milles, S., Jensen, M. R., Salvi, N., Kleman, J.-P., Maurin, D., Ruigrok, R. W. H., and Blackledge, M. (2020) Measles virus nucleo-and phosphoproteins form liquid-like phase-separated compartments that promote nucleocapsid assembly. Science advances 6, eaaz7095–eaaz7095

12. Zhou, Y., Su Justin, M., Samuel Charles, E., Ma, D., and Dutch Rebecca, E. Measles Virus Forms Inclusion Bodies with Properties of Liquid Organelles. J Virol 93, e00948–00919

13. Katoh, H., Kubota, T., Kita, S., Nakatsu, Y., Aoki, N., Mori, Y., Maenaka, K., Takeda, M., and Kidokoro, M. (2015) Heat shock protein 70 regulates degradation of the mumps virus phosphoprotein via the ubiquitin-proteasome pathway. J Virol 89, 3188–3199

14. Carlos, T. S., Young, D. F., Schneider, M., Simas, J. P., and Randall, R. E. (2009) Parainfluenza virus 5 genomes are located in viral cytoplasmic bodies whilst the virus dismantles the interferon-induced antiviral state of cells. J Gen Virol 90, 2147–2156

15. Zhang, S., Chen, L., Zhang, G., Yan, Q., Yang, X., Ding, B., Tang, Q., Sun, S., Hu, Z., and Chen, M. (2013) An amino acid of human parainfluenza virus type 3 nucleoprotein is critical for template function and cytoplasmic inclusion body formation. J Virol 87, 12457–12470

16. Ringel, M., Heiner, A., Behner, L., Halwe, S., Sauerhering, L., Becker, N., Dietzel, E., Sawatsky, B., Kolesnikova, L., and Maisner, A. (2019) Nipah virus induces two inclusion body populations: Identification of novel inclusions at the plasma membrane. PLoS Pathog 15, e1007733–e1007733

17. Patton, J. T., Silvestri, L. S., Tortorici, M. A., Vasquez-Del Carpio, R., and Taraporewala, Z. F. (2006) Rotavirus Genome Replication and Morphogenesis: Role of the Viroplasm. in Reoviruses: Entry, Assembly and Morphogenesis (Roy, P. ed.), Springer Berlin Heidelberg, Berlin, Heidelberg. pp 169–187

18. Silvestri, L. S., Taraporewala, Z. F., and Patton, J. T. (2004) Rotavirus replication: plus-sense templates for double-stranded RNA synthesis are made in viroplasms. J Virol 78, 7763–7774

19. Tenorio, R., Fernández de Castro, I., Knowlton, J. J., Zamora, P. F., Sutherland, D. M., Risco, C., and Dermody, T. S. (2019) Function, Architecture, and Biogenesis of Reovirus Replication Neoorganelles. Viruses 11, 288

20. Nevers, Q., Albertini, A. A., Lagaudrière-Gesbert, C., and Gaudin, Y. (2020) Negri bodies and other virus membrane-less replication compartments. Biochim Biophys Acta Mol Cell Res 1867, 118831–118831

21. Savastano, A., Ibáñez de Opakua, A., Rankovic, M., and Zweckstetter, M. (2020) Nucleocapsid protein of SARS-CoV-2 phase separates into RNA-rich polymerase-containing condensates. Nature Communications 11, 6041

22. Cong, Y., Ulasli, M., Schepers, H., Mauthe, M., V’Kovski, P., Kriegenburg, F., Thiel, V., de Haan, C. A. M., and Reggiori, F. (2020) Nucleocapsid Protein Recruitment to Replication-Transcription Complexes Plays a Crucial Role in Coronaviral Life Cycle. J Virol 94, e01925–01919

23. Iserman, C., Roden, C. A., Boerneke, M. A., Sealfon, R. S. G., McLaughlin, G. A., Jungreis, I., Fritch, E. J., Hou, Y. J., Ekena, J., Weidmann, C. A., Theesfeld, C. L., Kellis, M., Troyanskaya, O. G., Baric, R. S., Sheahan, T. P., Weeks, K. M., and Gladfelter, A. S. (2020) Genomic RNA Elements Drive Phase Separation of the SARS-CoV-2 Nucleocapsid. Molecular Cell 80, 1078–1091.e1076

24. Chen, H., Cui, Y., Han, X., Hu, W., Sun, M., Zhang, Y., Wang, P.-H., Song, G., Chen, W., and Lou, J. (2020) Liquid–liquid phase separation by SARS-CoV-2 nucleocapsid protein and RNA. Cell Res 30, 1143–1145

25. Cubuk, J., Alston, J. J., Incicco, J. J., Singh, S., Stuchell-Brereton, M. D., Ward, M. D., Zimmerman, M. I., Vithani, N., Griffith, D., Wagoner, J. A., Bowman, G. R., Hall, K. B., Soranno, A., and Holehouse, A. S. (2021) The SARS-CoV-2 nucleocapsid protein is dynamic, disordered, and phase separates with RNA. Nature communications 12, 1936–1936

26. Jack, A., Ferro, L. S., Trnka, M. J., Wehri, E., Nadgir, A., Nguyenla, X., Fox, D., Costa, K., Stanley, S., Schaletzky, J., and Yildiz, A. (2021) SARS-CoV-2 nucleocapsid protein forms condensates with viral genomic RNA. PLOS Biology 19, e3001425

27. Lu, S., Ye, Q., Singh, D., Cao, Y., Diedrich, J. K., Yates, J. R., Villa, E., Cleveland, D. W., and Corbett, K. D. (2021) The SARS-CoV-2 nucleocapsid phosphoprotein forms mutually exclusive condensates with RNA and the membrane-associated M protein. Nature Communications 12, 502

28. Perdikari, T. M., Murthy, A. C., Ryan, V. H., Watters, S., Naik, M. T., and Fawzi, N. L. (2020) SARS-CoV-2 nucleocapsid protein phase-separates with RNA and with human hnRNPs. EMBO J 39, e106478–e106478

29. Carlson, C. R., Asfaha, J. B., Ghent, C. M., Howard, C. J., Hartooni, N., Safari, M., Frankel, A. D., and Morgan, D. O. (2020) Phosphoregulation of Phase Separation by the SARS-CoV-2&#xa0;N Protein Suggests a Biophysical Basis for its Dual Functions. Molecular Cell 80, 1092–1103.e1094

30. Wheeler, R. J., Lee, H. O., Poser, I., Pal, A., Doeleman, T., Kishigami, S., Kour, S., Anderson, E. N., Marrone, L., Murthy, A. C., Jahnel, M., Zhang, X., Boczek, E., Fritsch, A., Fawzi, N. L., Sterneckert, J., Pandey, U., David, D. C., Davis, B. G., Baldwin, A. J., Hermann, A., Bickle, M., Alberti, S., and Hyman, A. A. (2019) Small molecules for modulating protein driven liquid-liquid phase separation in treating neurodegenerative disease. bioRxiv, 721001

31. Fang, M. Y., Markmiller, S., Vu, A. Q., Javaherian, A., Dowdle, W. E., Jolivet, P., Bushway, P. J., Castello, N. A., Baral, A., Chan, M. Y., Linsley, J. W., Linsley, D., Mercola, M., Finkbeiner, S., Lecuyer, E., Lewcock, J. W., and Yeo, G. W. (2019) Small-Molecule Modulation of TDP-43 Recruitment to Stress Granules Prevents Persistent TDP-43 Accumulation in ALS/FTD. Neuron 103, 802–819.e811

32. Risso-Ballester, J., Galloux, M., Cao, J., Le Goffic, R., Hontonnou, F., Jobart-Malfait, A., Desquesnes, A., Sake, S. M., Haid, S., Du, M., Zhang, X., Zhang, H., Wang, Z., Rincheval, V., Zhang, Y., Pietschmann, T., Eléouët, J.-F., Rameix-Welti, M.-A., and Altmeyer, R. (2021) A condensate-hardening drug blocks RSV replication in vivo. Nature 595, 596–599

33. Conti, B. A., and Oppikofer, M. (2022) Biomolecular condensates: new opportunities for drug discovery and RNA therapeutics. Trends in Pharmacological Sciences 43, 820–837

34. Mitrea, D. M., Mittasch, M., Gomes, B. F., Klein, I. A., and Murcko, M. A. (2022) Modulating biomolecular condensates: a novel approach to drug discovery. Nature Reviews Drug Discovery

35. Biesaga, M., Frigolé-Vivas, M., and Salvatella, X. (2021) Intrinsically disordered proteins and biomolecular condensates as drug targets. Curr Opin Chem Biol 62, 90–100

36. Cai, T., Yu, Z., Wang, Z., Liang, C., and Richard, S. (2021) Arginine methylation of SARS-Cov-2 nucleocapsid protein regulates RNA binding, its ability to suppress stress granule formation, and viral replication. J Biol Chem 297, 100821

37. Doble, B. W., Patel, S., Wood, G. A., Kockeritz, L. K., and Woodgett, J. R. (2007) Functional redundancy of GSK-3alpha and GSK-3beta in Wnt/beta-catenin signaling shown by using an allelic series of embryonic stem cell lines. Dev Cell 12, 957–971

38. Wu, C. H., Yeh, S. H., Tsay, Y. G., Shieh, Y. H., Kao, C. L., Chen, Y. S., Wang, S. H., Kuo, T. J., Chen, D. S., and Chen, P. J. (2009) Glycogen synthase kinase-3 regulates the phosphorylation of severe acute respiratory syndrome coronavirus nucleocapsid protein and viral replication. J Biol Chem 284, 5229–5239

39. Liu, X., Verma, A., Garcia, G., Ramage, H., Lucas, A., Myers, R. L., Michaelson, J. J., Coryell, W., Kumar, A., Charney, A. W., Kazanietz, M. G., Rader, D. J., Ritchie, M. D., Berrettini, W. H., Schultz, D. C., Cherry, S., Damoiseaux, R., Arumugaswami, V., and Klein, P. S. (2021) Targeting the coronavirus nucleocapsid protein through GSK-3 inhibition. Proceedings of the National Academy of Sciences 118, e2113401118

40. Rudd, C. E. (2020) GSK-3 Inhibition as a Therapeutic Approach Against SARs CoV2: Dual Benefit of Inhibiting Viral Replication While Potentiating the Immune Response. Front Immunol 11, 1638

41. Pillaiyar, T., and Laufer, S. (2022) Kinases as Potential Therapeutic Targets for Anti-coronaviral Therapy. Journal of Medicinal Chemistry 65, 955–982

42. Wu, D., and Pan, W. (2010) GSK3: a multifaceted kinase in Wnt signaling. Trends Biochem Sci 35, 161–168

43. Polakis, P. (1999) The oncogenic activation of β-catenin. Current Opinion in Genetics & Development 9, 15–21

44. Martínez-González, L., Gonzalo-Consuegra, C., Gómez-Almería, M., Porras, G., de Lago, E., Martín-Requero, Á., and Martínez, A. (2021) Tideglusib, a Non-ATP Competitive Inhibitor of GSK-3β as a Drug Candidate for the Treatment of Amyotrophic Lateral Sclerosis. Int J Mol Sci 22

45. Murer, L., Volle, R., Andriasyan, V., Petkidis, A., Gomez-Gonzalez, A., Yang, L., Meili, N., Suomalainen, M., Bauer, M., Policarpo Sequeira, D., Olszewski, D., Georgi, F., Kuttler, F., Turcatti, G., and Greber, U. F. (2022) Identification of broad anti-coronavirus chemical agents for repurposing against SARS-CoV-2 and variants of concern. Curr Res Virol Sci 3, 100019

46. Yang, L., Pei, R.-j., Li, H., Ma, X.-n., Zhou, Y., Zhu, F.-h., He, P.-l., Tang, W., Zhang, Y.-c., Xiong, J., Xiao, S.-q., Tong, X.-k., Zhang, B., and Zuo, J.-p. (2021) Identification of SARS-CoV-2 entry inhibitors among already approved drugs. Acta Pharmacologica Sinica 42, 1347–1353

47. Bouhaddou, M., Memon, D., Meyer, B., White, K. M., Rezelj, V. V., Correa Marrero, M., Polacco, B. J., Melnyk, J. E., Ulferts, S., Kaake, R. M., Batra, J., Richards, A. L., Stevenson, E., Gordon, D. E., Rojc, A., Obernier, K., Fabius, J. M., Soucheray, M., Miorin, L., Moreno, E., Koh, C., Tran, Q. D., Hardy, A., Robinot, R., Vallet, T., Nilsson-Payant, B. E., Hernandez-Armenta, C., Dunham, A., Weigang, S., Knerr, J., Modak, M., Quintero, D., Zhou, Y., Dugourd, A., Valdeolivas, A., Patil, T., Li, Q., Hüttenhain, R., Cakir, M., Muralidharan, M., Kim, M., Jang, G., Tutuncuoglu, B., Hiatt, J., Guo, J. Z., Xu, J., Bouhaddou, S., Mathy, C. J. P., Gaulton, A., Manners, E. J., Félix, E., Shi, Y., Goff, M., Lim, J. K., McBride, T., O’Neal, M. C., Cai, Y., Chang, J. C. J., Broadhurst, D. J., Klippsten, S., De Wit, E., Leach, A. R., Kortemme, T., Shoichet, B., Ott, M., Saez-Rodriguez, J., tenOever, B. R., Mullins, R. D., Fischer, E. R., Kochs, G., Grosse, R., García-Sastre, A., Vignuzzi, M., Johnson, J. R., Shokat, K. M., Swaney, D. L., Beltrao, P., and Krogan, N. J. (2020) The Global Phosphorylation Landscape of SARS-CoV-2 Infection. Cell 182, 685–712.e619

48. Patten, J. J., Keiser, P. T., Gysi, D., Menichetti, G., Mori, H., Donahue, C. J., Gan, X., Do Valle, I., Geoghegan-Barek, K., Anantpadma, M., Berrigan, J. L., Jalloh, S., Ayazika, T., Wagner, F., Zitnik, M., Ayehunie, S., Anderson, D., Loscalzo, J., Gummuluru, S., Namchuk, M. N., Barabasi, A. L., and Davey, R. A. (2021) Multidose evaluation of 6,710 drug repurposing library identifies potent SARS-CoV-2 infection inhibitors &lt;em&gt;In Vitro&lt;/em&gt; and &lt;em&gt;In Vivo&lt;/em&gt. bioRxiv, 2021.2004.2020.440626

49. Xiao, X., Wang, C., Chang, D., Wang, Y., Dong, X., Jiao, T., Zhao, Z., Ren, L., Dela Cruz, C. S., Sharma, L., Lei, X., and Wang, J. (2020) Identification of Potent and Safe Antiviral Therapeutic Candidates Against SARS-CoV-2. Frontiers in Immunology 11

50. Shen, Z., Halberg, A., Fong, J. Y., Guo, J., Song, G., Louie, B., Luedtke, G. R., Visuthikraisee, V., Protter, A., Koh, X., Baik, T., and Lum, P. Y. (2022) Elucidating host cell response pathways and repurposing therapeutics for SARS-CoV-2 and other coronaviruses using gene expression profiles of chemical and genetic perturbations. bioRxiv, 2022.2004.2018.488682

51. Beurel, E., Grieco, S. F., and Jope, R. S. (2015) Glycogen synthase kinase-3 (GSK3): regulation, actions, and diseases. Pharmacol Ther 148, 114–131

52. Kilgore, H. R., and Young, R. A. (2022) Learning the chemical grammar of biomolecular condensates. Nat Chem Biol

53. Nabeel-Shah, S., Lee, H., Ahmed, N., Burke, G. L., Farhangmehr, S., Ashraf, K., Pu, S., Braunschweig, U., Zhong, G., Wei, H., Tang, H., Yang, J., Marcon, E., Blencowe, B. J., Zhang, Z., and Greenblatt, J. F. (2022) SARS-CoV-2 nucleocapsid protein binds host mRNAs and attenuates stress granules to impair host stress response. iScience 25, 103562

54. Zhao, M., Yu, Y., Sun, L.-M., Xing, J.-Q., Li, T., Zhu, Y., Wang, M., Yu, Y., Xue, W., Xia, T., Cai, H., Han, Q.-Y., Yin, X., Li, W.-H., Li, A.-L., Cui, J., Yuan, Z., Zhang, R., Zhou, T., Zhang, X.-M., and Li, T. (2021) GCG inhibits SARS-CoV-2 replication by disrupting the liquid phase condensation of its nucleocapsid protein. Nature Communications 12, 2114

55. Zhang, N., and Ashizawa, T. (2017) RNA toxicity and foci formation in microsatellite expansion diseases. Current Opinion in Genetics & Development 44, 17–29

56. Shelkovnikova, T. A., Robinson, H. K., Southcombe, J. A., Ninkina, N., and Buchman, V. L. (2014) Multistep process of FUS aggregation in the cell cytoplasm involves RNA-dependent and RNA-independent mechanisms. Human Molecular Genetics 23, 5211–5226

57. Kino, Y., Washizu, C., Aquilanti, E., Okuno, M., Kurosawa, M., Yamada, M., Doi, H., and Nukina, N. (2011) Intracellular localization and splicing regulation of FUS/TLS are variably affected by amyotrophic lateral sclerosis-linked mutations. Nucleic Acids Research 39, 2781–2798

58. Takanashi, K., and Yamaguchi, A. (2014) Aggregation of ALS-linked FUS mutant sequesters RNA binding proteins and impairs RNA granules formation. Biochemical and Biophysical Research Communications 452, 600–607

59. Rodriguez, S., Sahin, A., Schrank, B. R., Al-Lawati, H., Costantino, I., Benz, E., Fard, D., Albers, A. D., Cao, L., Gomez, A. C., Evans, K., Ratti, E., Cudkowicz, M., Frosch, M. P., Talkowski, M., Sorger, P. K., Hyman, B. T., and Albers, M. W. (2021) Genome-encoded cytoplasmic double-stranded RNAs, found in C9ORF72 ALS-FTD brain, propagate neuronal loss. Science Translational Medicine 13, eaaz4699

60. Beausoleil, S. A., Villén, J., Gerber, S. A., Rush, J., and Gygi, S. P. (2006) A probability-based approach for high-throughput protein phosphorylation analysis and site localization. Nature Biotechnology 24, 1285–1292

61. Sanjana, N. E., Shalem, O., and Zhang, F. (2014) Improved vectors and genome-wide libraries for CRISPR screening. Nat Methods 11, 783–784

62. Doench, J. G., Fusi, N., Sullender, M., Hegde, M., Vaimberg, E. W., Donovan, K. F., Smith, I., Tothova, Z., Wilen, C., Orchard, R., Virgin, H. W., Listgarten, J., and Root, D. E. (2016) Optimized sgRNA design to maximize activity and minimize off-target effects of CRISPR-Cas9. Nat Biotechnol 34, 184–191

63. Xu, Q., Wang, Y., Dabdoub, A., Smallwood, P. M., Williams, J., Woods, C., Kelley, M. W., Jiang, L., Tasman, W., Zhang, K., and Nathans, J. (2004) Vascular development in the retina and inner ear: control by Norrin and Frizzled-4, a high-affinity ligand-receptor pair. Cell 116, 883–895

