## Supplementary Materials for "A screen for modulation of nucleocapsid protein condensation identifies small molecules with anti-coronavirus activity"

**SUPPLEMENTARY FIGURE 1 – Properties of HCoV N condensates.**

(A) Western blot quantification of A549 cells stably expressing the seven different N-EGFP proteins. Blots were stained with an anti-GFP antibody.

(B) Fluorescence images of a stable A549 EGFP cell line treated with 1µg/mL transfected polyIC for 7h. Images taken at 40x air magnification, confocal. Scale bar, 20 µm.

(C) Time-lapse imaging showing polyIC-induced and constitutive N condensate fusion events over time. Fusion events are identified by a dotted circle. Images taken at 40x air magnification, confocal. Scale bar, 10 µm.

(D) FRAP analyses of polyIC-induced and constitutive N condensates. Images are single N condensates representative of each condition. Mean fluorescence intensity plot illustrates FRAP results for n = 7 condensates per condition. Images taken at 60x oil magnification, widefield. Scale bar, 1 µm.

(E) Fluorescence images and quantification of HEK293T cells transiently transfected with various 'domain-swap' mutants of SARS-CoV-2 N, as indicated by the schematic. Data indicates mean ± standard deviation of duplicate experiments. Images taken at 40x air magnification, widefield. Scale bar, 10 µm.

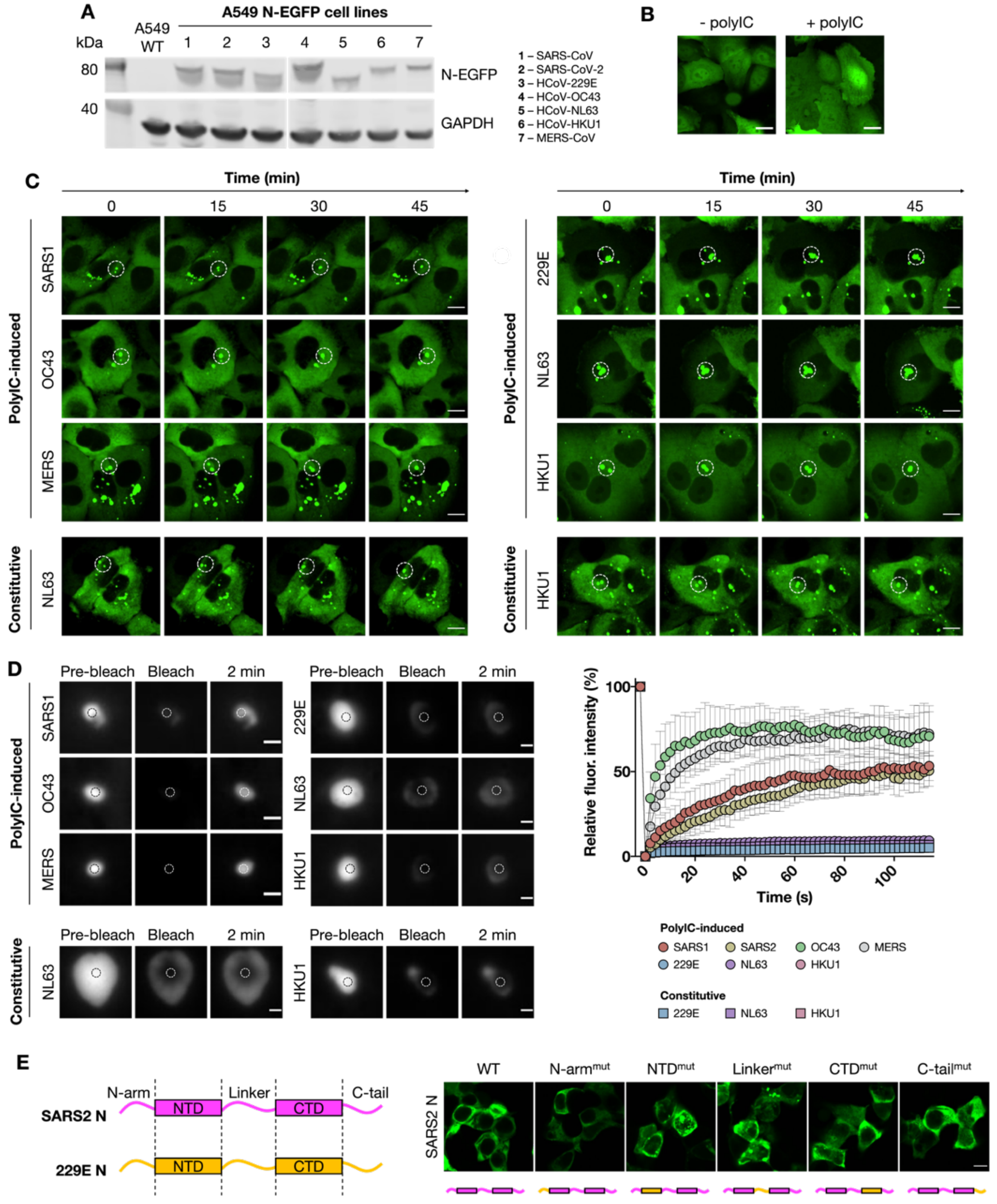

**SUPPLEMENTARY FIGURE 2 – High content screening and automated image analysis pipelines**

(A) Screening and automated image analysis pipeline for the primary screens performed. Experimental set-up (left panel), image acquisition and analysis with the SpotDetector.V4 algorithm (middle panel) and screening data output (right panel) are illustrated. Data output is represented as percentage N condensation or condensate inhibition activity.

(B) Screening follow up and automated image analysis pipeline for hit follow up experiments. Experimental set-up (left panel), image acquisition and analysis with a custom MetaXpress analysis pipeline (middle panel) and dose response data output (right panel) are illustrated.

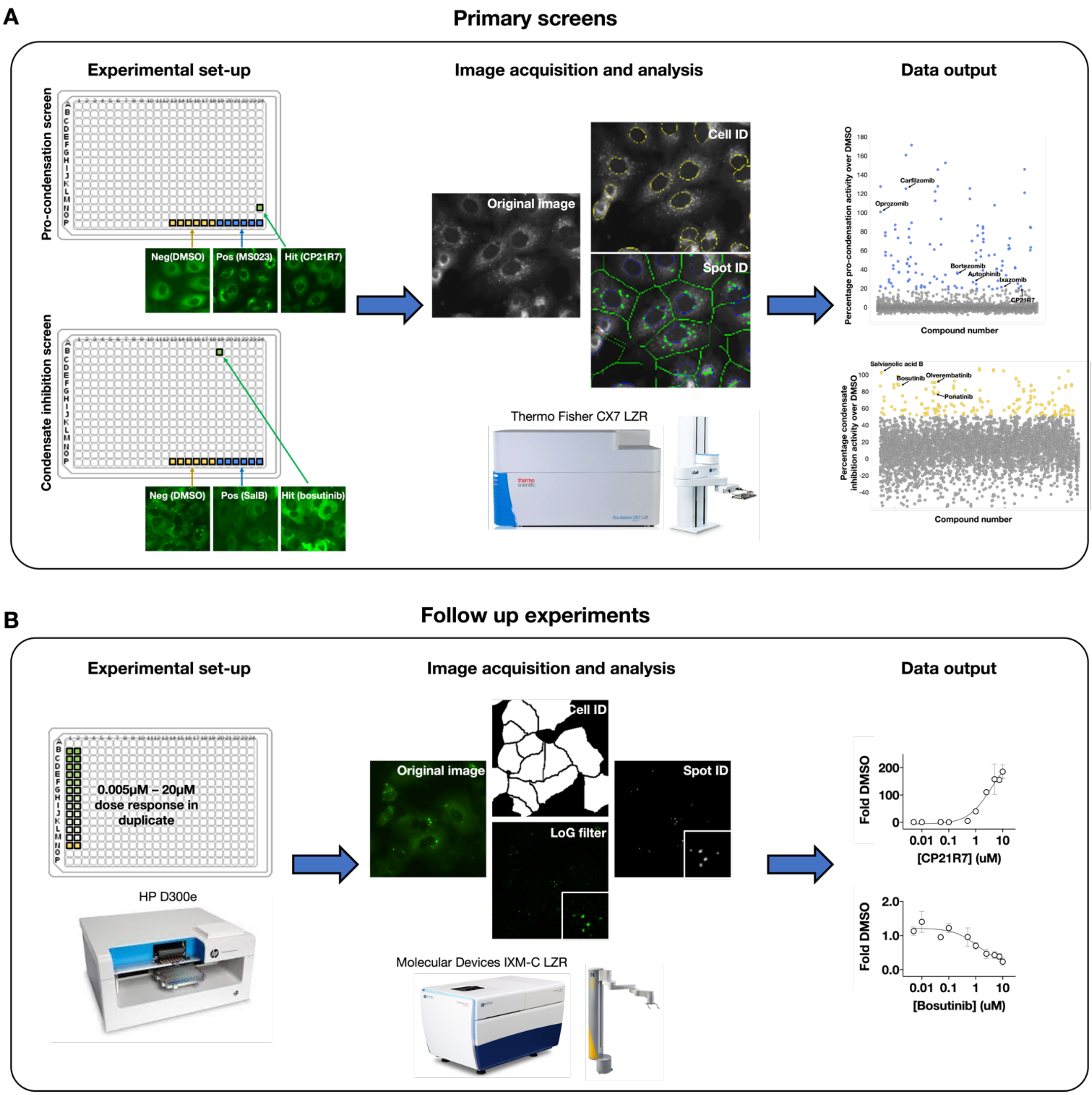

**SUPPLEMENTARY FIGURE 3 – ATP-competitive GSK3 and proteasome inhibitors induce SARS-CoV-2 N condensation.**

(A) Dose response curves illustrating increased nuclear localization of N upon treatment with the proteasome inhibitors. Curves indicate the fold change in average nuclear-to-cytoplasmic ratio of N intensity over DMSO control and illustrate mean ± standard deviation of duplicates.

(B) Fluorescence images of A549 SARS-CoV-2 N-EGFP cell line treated 10μM small molecules and 50mM LiCl for 24h. Images taken at 40x air magnification, widefield. Scale bar, 10 μm.

(C) Dose response curves for SARS-CoV-2 N condensation upon treatment with 10μM small molecules. Curves indicate fold change in number of N puncta per cell over DMSO control (fold DMSO) and illustrate mean ± standard deviation of duplicates.

(D) Mass spectrometric analyses data illustrating inhibition of phosphorylation at various phosphosites across the SARS-CoV-2 N protein by 1μM LY2090314 and 1μM CP21R7 treatment. Schematic illustrates locations of phosphopeptides identified on SARS-CoV-2 N. Exact phosphorylation sites, where confidently assigned, are identified in red.

(E) Fluorescence images of HEK293T cells transiently transfected with SARS-CoV-2 N-EGFP (WT N) and SARS-CoV-2 $N^{SAmut}$-EGFP. Images taken at 20x air magnification, widefield. WT: wild type. Scale bar, 20 μm.

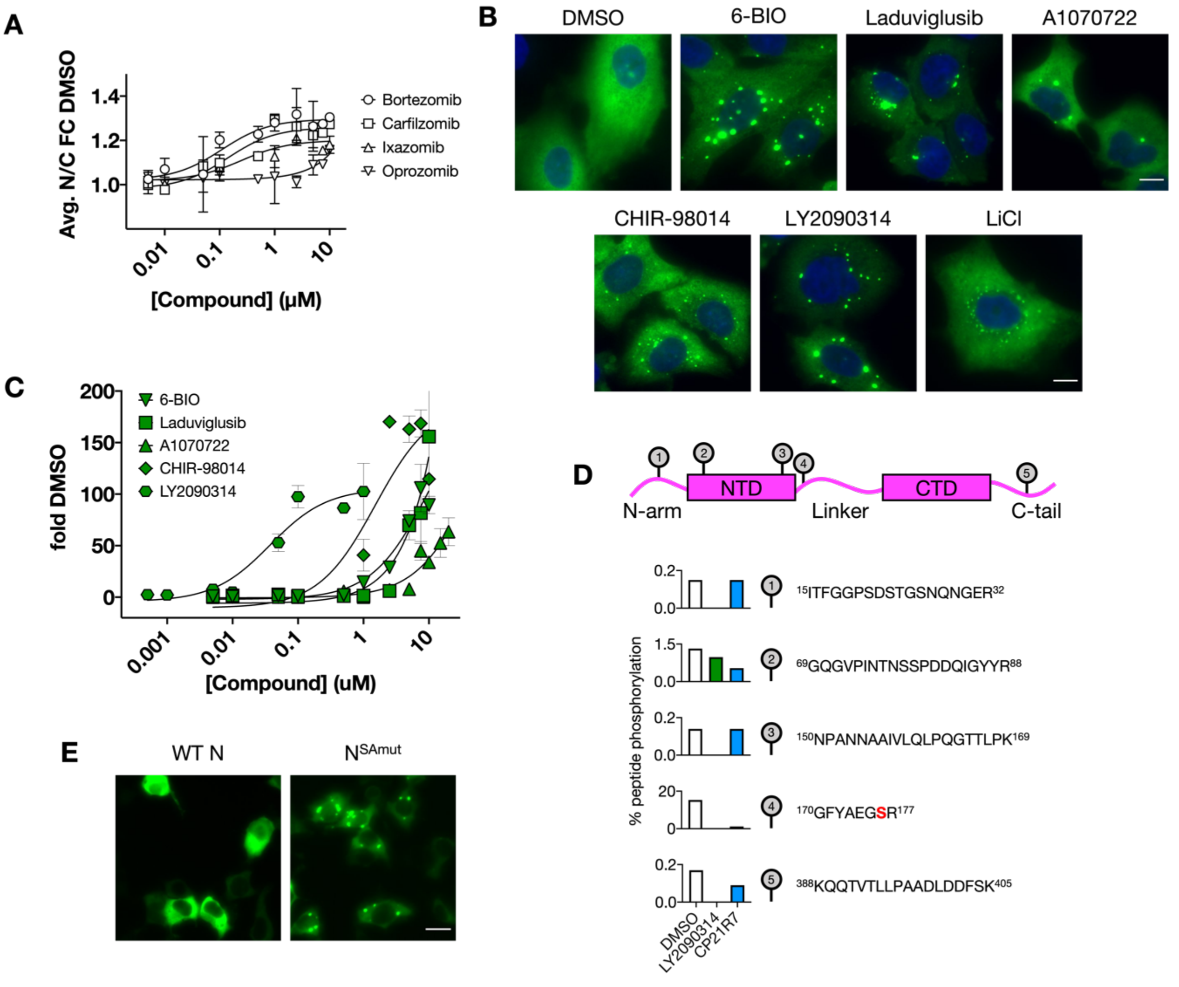

**SUPPLEMENTARY FIGURE 4 – ATP-competitive GSK3 inhibitors exhibit pan-HCoV N condensation activity and induce N condensate hardening.**

(A) Representative fluorescence images of all seven HCoV N-EGFP cell lines treated with 10μM GSK3 inhibitors for 24h. Images are taken at 40x air magnification, widefield. Scale bar, 10 μm.

(B) Dose response curves for N condensation across all seven HCoV N-expressing A549 cell lines upon treatment with GSK3 inhibitors. Curves indicate fold change in number of N puncta per cell over DMSO control (fold DMSO) and illustrate mean ± standard deviation of duplicates.

(C) Fluorescence images and quantification of HEK293T cells transiently transfected with various bat-CoV N-EGFP and treated with 0.1μM LY2090314. Images are taken at 20x air magnification, confocal. Scale bar, 20 μm.

(D) FRAP analyses of SARS-CoV, SARS-CoV-2, HCoV-OC43 and MERS-CoV polyIC-induced and LY2090314-induced condensates. Mean fluorescence intensity plot illustrates FRAP results for n = 7 condensates per condition.

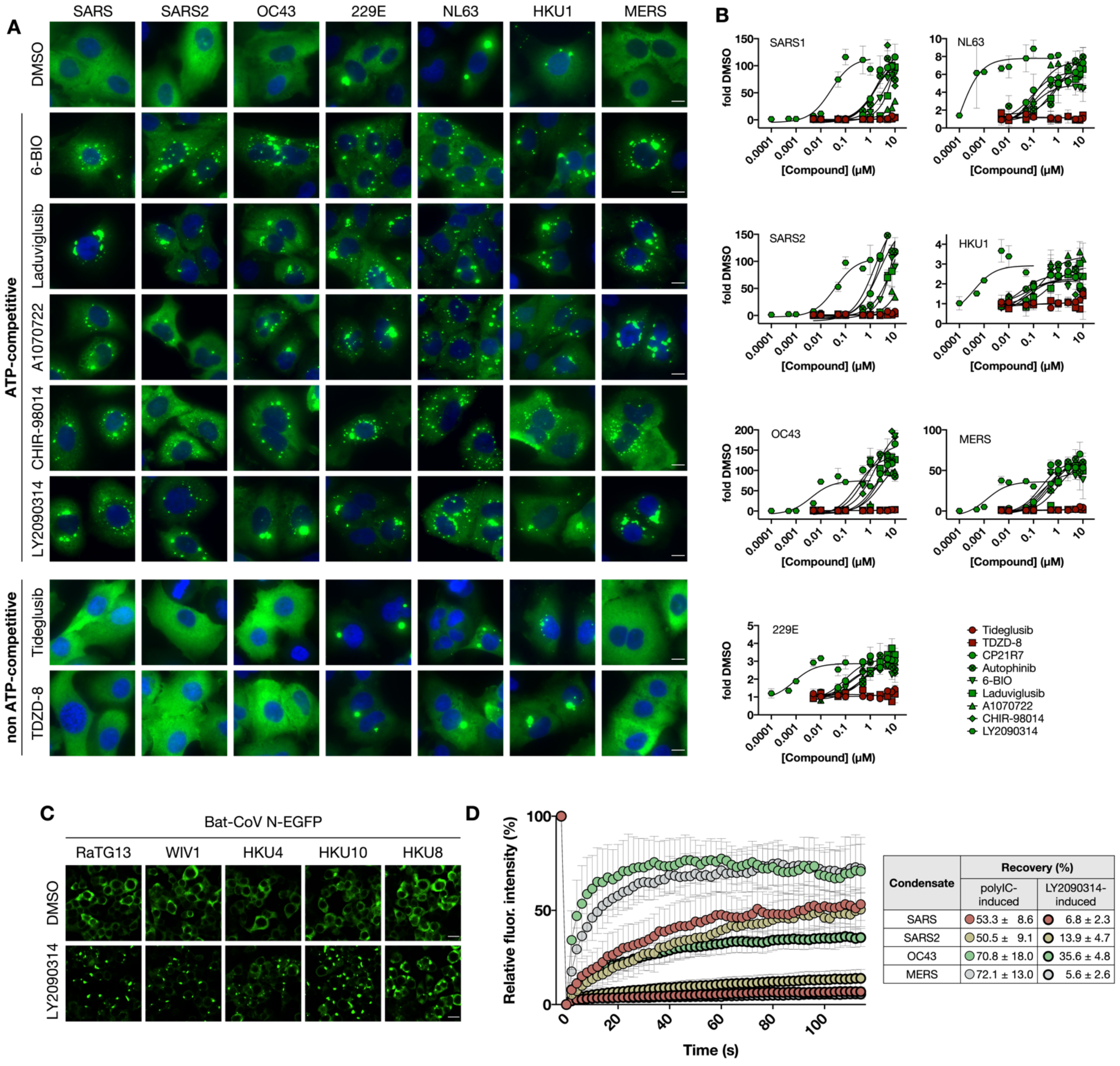


| Condensate | Recovery (%) | |
|---|---|---|
| | polyIC-induced | LY2090314-induced |
| SARS | 53.3 ± 8.6 | 6.8 ± 2.3 |
| SARS2 | 50.5 ± 9.1 | 13.9 ± 4.7 |
| OC43 | 70.8 ± 18.0 | 35.6 ± 4.8 |
| MERS | 72.1 ± 13.0 | 5.6 ± 2.6 |

**SUPPLEMENTARY FIGURE 5 – Condensate inhibitors block the polyIC input and exhibit multi-HCoV condensation inhibition activity.**

(A) Fluorescence images illustrating inhibition of IRF3 activation and stress granule formation by various N condensate inhibitors, as represented by a decrease in nuclear localization of IRF3 or a decrease in number of G3BP1 puncta, respectively. Images are taken at 40x air magnification, widefield. Scale bar, 50 µm.

(B) Fluorescence images of A549 SARS-CoV-2 N-EGFP cell line treated 10µM imatinib, saracatinib, PP2 or 7.5µM nilotinib for 17h followed by 1µg/mL transfected polyIC treatment for a further 7h. Images taken at 40x air magnification, widefield. Dose response curves indicate fold change of number of N puncta per cell over DMSO control (fold DMSO) and illustrate mean ± standard deviation of duplicates. Scale bar, 10 µm.

(C) Dose response curves for N condensation across SARS-CoV, SARS-CoV-2, HCoV-OC43 and MERS-CoV N-expressing A549 cell lines upon treatment with condensate inhibitors for 17h followed by 1µg/mL transfected polyIC treatment for a further 7h. Curves indicate fold change in number of N puncta per cell over DMSO control (fold DMSO) and illustrate mean ± standard deviation of duplicates.

(D) Dose response curves for N condensation across HCoV-229E, HCoV-NL63 and HCoV-HKU1 N-expressing A549 cell lines upon treatment with condensate inhibitors, without the addition of polyIC. Curves indicate fold change in number of N puncta per cell over DMSO control (fold DMSO) and illustrate mean ± standard deviation of duplicates.

(E) Dose response curves for N condensation across HCoV-229E, HCoV-NL63 and HCoV-HKU1 N-expressing A549 cell lines upon treatment with condensate inhibitors for 17h followed by 1µg/mL transfected polyIC treatment for a further 7h. Curves indicate fold change in number of N puncta per cell over DMSO control (fold DMSO) and illustrate mean ± standard deviation of duplicates.

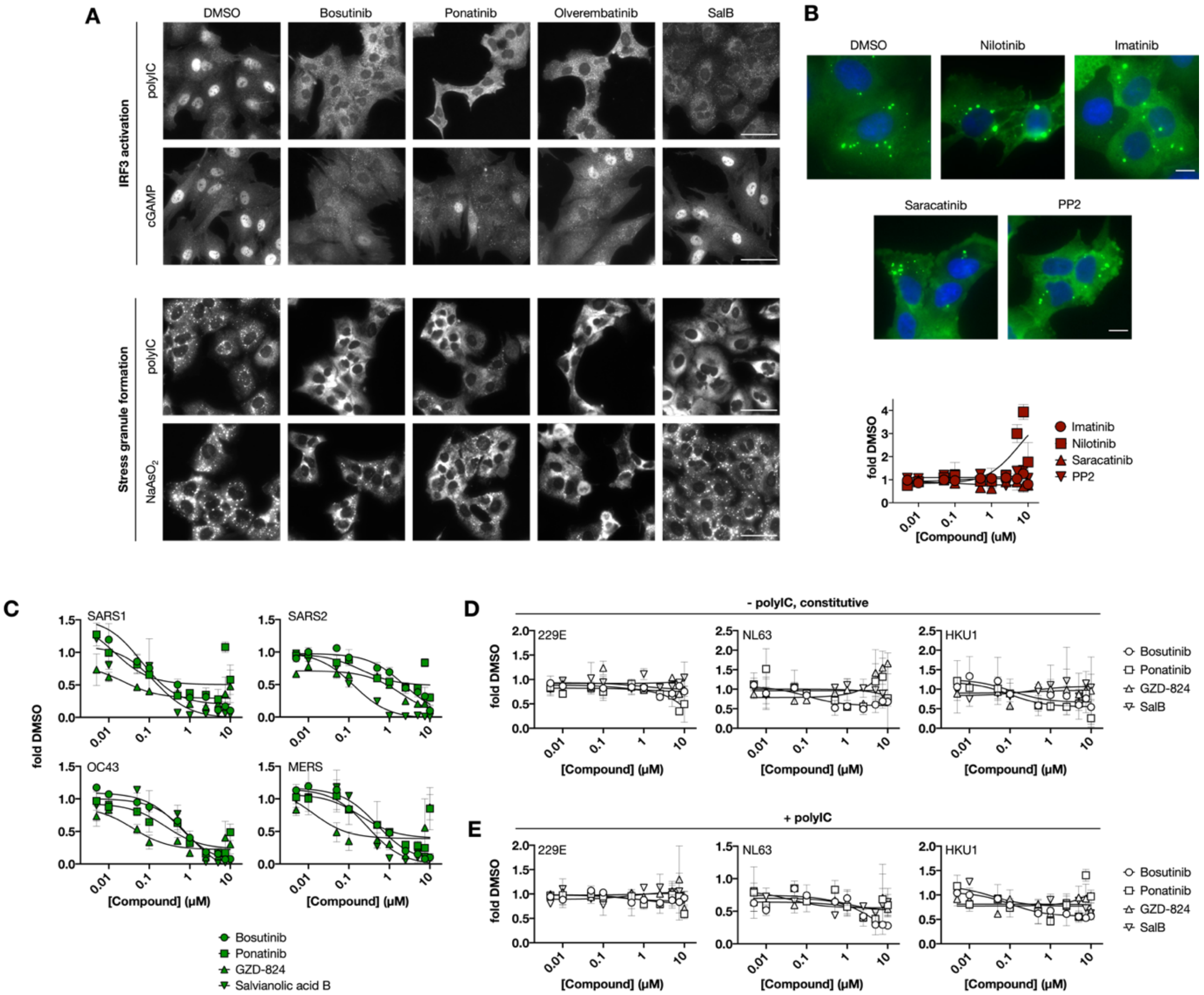

**SUPPLEMENTARY TABLE 1 – Small molecule screening data**

| Category | Parameter | Description |
|---|---|---|
| Assay | Type of assay | Cell-based |
| | Target | SARS-CoV-2 N fused to a C-terminal EGFP |
| | Primary measurement | Fluorescence imaging, image analysis and quantification |
| | Key reagents | A549 SARS-CoV-2 N-EGFP cell line, low molecular weight polyIC (InVivoGen tlrl-picw), Lipofectamine3000™ (Thermo Fisher), Selleck Chemicals FDA-approved compound library comprising 2,554 compounds (Selleck Chemicals), 10% methanol-free formaldehyde diluted to 3% in PBS, Hoechst 33342 (Thermo Scientific 62249) |
| | Assay protocol | A small molecule library was screened against the A549 SARS-CoV-2 N-EGFP cell line to identify small molecules that either promote or inhibit N condensation (pro-condenser and condensate inhibitor screens respectively). The screen was performed at AbbVie. Briefly, for the pro-condensation screen, 2,000 cells were seeded in each well of a 384 well plate in a final volume of 30uL. The next day, cells were treated with compounds at a final concentration of 10µM for 24h. For the condensate inhibition screen, cells were additionally treated with 1µg/mL transfected polyIC 17h after compound addition, 7h before fixation. Cells were then fixed for 20min in 10% methanol-free formaldehyde diluted to 3% in PBS supplemented with 1µg/mL Hoechst 33342 and then washed with Biotek EL405 platewasher. Plates were then imaged with the Thermo Fisher CX7 LZR using a 20x objective and analyzed with the SpotDetector.V4 algorithm. The output feature SpotCountPerObject was used to evaluate the changes in N condensation in both screens. Compounds that showed activity greater than 2 standard deviations from the mean based on control percentage activity (pro-condensation or condensate inhibition) were selected. All assays were performed in duplicate. Hits were then validated with dose responses both at AbbVie and at Harvard Medical School with independent image analysis pipelines. |
| | Additional comments | - |
| Library | Library size | 2,554 |
| | Library composition | 2,082 FDA-approved drug library and 472 additional bioactives |
| | Source | Selleck Chemicals LLC |
| | Additional comments | - |
| Screen | Format | Perkin Elmer LLC ViewPlate-384 Black, Optically Clear Bottom, Tissue Culture Treated (PerkinElmer 6007460) |

| | | |
|---|---|---|
| | Concentration(s) tested | 10μM (0.1% final concentration of DMSO) |
| | Plate controls | Negative control: DMSO only<br>Positive control (pro-condensation screen): 10μM MS023)<br>Positive control (condensate inhibition screen): 10μM salvianolic acid B |
| | Reagent/ compound dispensing system | Primary screen: Thermo Scientific Multidrop™ Combi Reagent Dispenser (cells and polyIC dispensing), Labcyte Echo 555 (compound dispensing), Biomek FX (compound dilution, addition)<br>Follow up experiments: Thermo Scientific Multidrop™ Combi Reagent Dispenser (cells and polyIC dispensing), HP D300e Digitial Dispenser (compound dispensing) |
| | Detection instrument and software | Primary screen: Thermo Fisher CX7 LZR, HCS Studio software, SpotDetector.V4 algorithm<br>Follow up experiments: Molecular Devices ImageXpress Micro Confocal Laser (fluorescence microscope), MetaXpress High Content Image Acquisition & Analysis Software Version 62.7 (image analysis) |
| | Assay validation/QC | Z' factor (pro-condensation screen): 0.61<br>Z' factor (condensate inhibition screen): 0.55 |
| | Correction factors | - |
| | Normalization | Normalized to number of N puncta per cell based on % activity of controls, where % activity = (cMax-experimental/cMax-cMin) x 100, cMax = MS023 (pro-condensation screen) or DMSO (condensate inhibition screen) and cMin = DMSO (pro-condensation screen) or salvianolic acid B (condensate inhibition screen) |
| | Additional comments | - |
| Post-HTS analysis | Hit criteria | Hits were identified as compounds that showed pro-condensation or condensate inhibition activity greater than two standard deviations from the mean compared to DMSO control, after manual inspection and removal of artifacts |
| | Hit rate | Pro-condensation screen: 6/2,554 (0.23%)<br>Condensate inhibition screen: 4/2,554 (0.16%) |
| | Additional assay(s) | Counter-screens of hits against A549 wild-type cells (to identify and remove auto-fluorescent artifacts) and A549 EGFP cells (to identify and remove hits perturbing EGFP instead of N) |
| | Confirmation of hit purity and structure | Hit compounds were repurchased from MedChemExpress and re-tested in dose response format to confirm activity. |
| | Additional comments | - |
